# The influence of adaptation to life at high-altitude on condition dependent sexual shape and size dimorphism in *Drosophila melanogaster*

**DOI:** 10.1101/2021.06.30.450630

**Authors:** Maria Pesevski, Ian Dworkin

## Abstract

Sexual dimorphism is common despite factors such as inter-sex genetic correlations and sex-specific patterns of selection that might limit its evolution. Sexual dimorphism can be phenotypically plastic and condition dependent, that themselves may be targets of selection. However, it remains unclear how sexual dimorphism, its plasticity and condition dependence evolves, in particular during rapid adaptation to a new environment. Furthermore, the interplay between SSD and other forms of dimorphism, such as shape dimorphism co-evolves. Using Sub-Saharan populations of *Drosophila melanogaster* that vary for size and shape as a result of adaptation to high altitude environments, we examined sex specific patterns of developmental plasticity. We raised strains of *Drosophila* from low (Zambia) and high (Ethiopia) altitude populations varying for food quality or rearing temperature. We observed expected differences in wing size and shape due to population, sex and plasticity. While larval mass showed substantial evolved changes for sex specific condition dependence, effects on wing size and shape were modest. We examined shape-size allometric effects between groups. Allometric effects were generally similar across sexes, but differed substantially due to population of origin and plasticity. We discuss findings within the context of the evolution of plasticity for SSD, condition dependence and allometric relationships.

## Introduction

Sexual dimorphism is common and extremely captivating. While many studies examine species with male-biased sexual size dimorphism, particularly with exaggerated ornaments or weapons, female-biased SSD is more common, and with exceptions, (e.g. angler fish) differences are often subtle (Fairbairn, 2007). Sexual selection is frequently invoked to explain male-biased SSD, while fecundity selection seems to be a common explanation for female-biased SSD (Darwin, 1896; Shine, 1988; Pincheira-Donoso and Hunt, 2017) along with other ecological mechanisms (Hedrick and Temeles, 1989; Darwin, 1896; Selander, 1966; Shine, 1989). The study of the evolutionary origins and maintenance of sexual dimorphism remains an active area of interest for at least two reasons. First, despite potential antagonistic effects due to intra-locus sexual conflict and constraints due to high inter-sex genetic correlations, *r_fm_* (Poissant et al., 2010), sexual dimorphism occurs widely and may represent evolutionary resolution of conflict (Lande, 1980; Bonduriansky and Chenoweth, 2009; Cox and Calsbeek, 2009). Despite often high *r_fm_*, sexual dimorphism evolves rapidly in some contexts (Reeve and Fairbairn, 1999; Bird and Schaffer, 1972; Eisen and Hanrahan, 1972; Alicchio and Palenzona, 1971) though not all (Stewart and Rice, 2018; Tigreros and Lewis, 2011; Reeve and Fairbairn, 1996). Second, sexually dimorphic traits often are multi-functional and reflect the balance between sex-specific and shared functions that are targets of selection, potentially influencing multiple fitness components (Lande, 1980; Lasne et al., 2018; Matthews et al., 2019; Connallon, 2010; Connallon and Hall, 2016; Connallon et al., 2018).

It is important to consider how adaptive evolution on such multi-functional traits influences sexual dimorphism (Cox and Calsbeek, 2009) and how sexual dimorphism and high *r_fm_* influence trait evolution towards new adaptive optima (Lande, 1980; Connallon, 2015). Adaptation to a new environment may be influenced by sexually antagonistic selection and *r_fm_* (Lande, 1980; Cox and Calsbeek, 2009). When selection is consistent across sexes in the new environment, a generally high *r_fm_* may facilitate the adaptive process, though the process may be limited in the presence of sexually antagonistic selection (Connallon, 2015; Connallon and Hall, 2016). If selection is discordant between the sexes (Cox and Calsbeek, 2009), with respect to the new environment, this can constrain adaptive responses. For instance, the “gender load” imposed by sex specific effects on rate of adaptation has been estimated to be on the order of 50% in a wild population of *Parus major* (Poissant et al., 2016). However, the degree to which selection is generally discordant is unclear (Singh and Punzalan, 2018), and issues such as changes in *r_fm_* across environments (Punzalan et al., 2014) and imprecision in estimates of selection (Morrissey, 2016) makes this a challenging empirical and theoretical undertaking.

Studies of sexual dimorphism for morphology often focus on sexual size dimorphism (SSD), often examining a single trait at a time. However, a multivariate perspective in general, and sexual shape dimorphism (SShD) in particular is increasingly recognized as both pervasive and important (Gidaszewski et al., 2009; Abbott et al., 2010; Sztepanacz and Houle, 2019, 2021). Using multivariate (Butler and Losos, 2002) and geometric morpho-metric approaches not only have increased sensitivity to detect subtle changes, but SShD itself appears to be a target of selection in some instances (Gidaszewski et al., 2009; Abbott et al., 2010; Menezes et al., 2013). In a multivariate context, sex specific patterns of selection and the influence of the inter-sex covariance matrix **B** can result in more complex patterns of response to selection (Wyman et al., 2013; Sztepanacz and Houle, 2019; Gosden et al., 2012; Poissant et al., 2016). Despite this, sexual shape dimorphism is clearly evolving in numerous systems (Houle et al., 2017; Gidaszewski et al., 2009; Sztepanacz and Houle, 2021; Sanger et al., 2013; Evans et al., 2019; Chazot et al., 2016). However allometric relationships between shape and size can contribute substantially to SShD (Butler and Losos, 2002; Kaliontzopoulou et al., 2008; Gidaszewski et al., 2009; Sztepanacz and Houle, 2021), making it difficult to disentangle direct and indirect effects. When both shape-size allometry and SSD occur, this will generate SShD, which can be complicated by sex-specific allometric relationships. SShD can also be generated by sex differences in covariance patterns for the structures contributing to shape. Partitioning the relative contributions of these factors and the degree to which they are evolving independently remains an important and ongoing research area into the evolution of SShD (Fernández-Montraveta and Marugan-Lobon, 2017; Kaliontzopoulou et al., 2008; Butler and Losos, 2002).

Further complicating the study of SSD and SShD is that size and shape are often phenotypically plastic (David et al., 2009; Bitner–Mathe and Klaczko, 1999; Imasheva et al., 1999; Karan et al., 2000; Bubliy et al., 2001; Bubliy and Loeschcke, 2002; Debat et al., 2003, 2009; Shingleton et al., 2009). Patterns of plasticity are often dependent on traits under consideration and environmental variables generating the plastic response (Shingleton et al., 2009). Plasticity for size may in turn influence patterns of trait covariation and allometry with shape. Nutritional plasticity is a particularly interesting case with regards to sexual dimorphism, as it is directly linked to the concept of organismal condition (Andersson, 1986; Iwasa et al., 1991; Rowe and Houle, 1996; Cotton et al., 2004b; Bonduriansky, 2007a,b). Condition reflects the contribution of both genetic (efficiency of resource utilization) and environmental (access to resources) factors (Cotton et al., 2004b; Hill, 2011). As such the most common experimental manipulation to study condition dependence has been to vary access to resources (Cotton et al., 2004b). While there are several hypotheses regarding the evolution of sex-specific plasticity (adaptive canalization and condition dependence chief amongst them), considerable evidence has demonstrated that the most sexually dimorphic tend to also be the most condition dependent (Andersson, 1986; Rowe and Houle, 1996; Bonduriansky, 2007a,b), a pattern which seems to also hold for sex-biased gene expression (Wyman et al., 2010; Zinna et al., 2018). Evidence supports that condition dependence is correlated with sexual dimorphism, in particular in male-biased SSD systems with highly exaggerated secondary sexual traits (Bonduriansky and Rowe, 2005; Bonduriansky, 2007a, 2009; Cotton et al., 2004a,b; Rohner and Blanckenhorn, 2018; Rohner et al., 2018b; Stillwell and Fox, 2007). However, there is considerably less theoretical and empirical work that explores the interplay between sexual dimorphism and condition when it comes to moderate female biased SSD, the most common type of sexual dimorphism in animals (Rohner and Blanckenhorn, 2018; Oudin et al., 2015). With some notable exceptions (Ceballos and Valenzuela, 2011), we still know relatively little about sex-specific plasticity for shape and SShD.

With many factors influencing trait variation, it is important to consider how strong directional selection on a trait resulting in local adaptation may result in correlated effects on sex-specific trait plasticity generally, and condition dependence more specifically. The evolution of *Drosophila* wing morphology is an excellent system to explore these questions. While there is extensive genetic and mutational variation for wing size and shape within and between populations (Mezey and Houle, 2005; Pitchers et al., 2013), it generally has very high *r_fm_* (Sztepanacz and Houle, 2019) and sex-concordant effects of mutations (Testa and Dworkin, 2016). Comparative studies suggest that *Drosophila* wing shape, and dimorphism for shape has evolved relatively slowly (Gidaszewski et al., 2009; Houle et al., 2017; Sztepanacz and Houle, 2021), yet there is substantial variation for SSD (Rohner et al., 2018a; Huey et al., 2006; Blanckenhorn et al., 2007; Gidaszewski et al., 2009) and SShD (Gidaszewski et al., 2009). Importantly, allometric effects have been implicated in accounting for close to 50% of the variation in SShD among species (Gidaszewski et al., 2009; Sztepanacz and Houle, 2021), although the consequences of sex-specific allometry have not been investigated thoroughly. In this study, we examine developmental plasticity in wing morphology in response to temperature and food quality manipulations (to specifically assess condition dependence) in a pair of populations of *Drosophila melanogaster* originating from different altitudes. Within *Drosophila melanogaster*, there is considerable evidence for intraspecific divergence along altitudinal and latitudinal clines in both its native and more recently colonized worldwide populations (clinal variation in Americas, Australia) (Gibert et al., 2004; James et al., 1997; David and Bocquet, 1975; Gilchrist et al., 2000; Hoffmann and Weeks, 2007; Azevedo et al., 1998). Wing morphology is a particularly interesting model system as it shows considerable plasticity in particular to nutrition, rearing temperature and oxygen partial pressure (David et al., 2009; Bitner–Mathe and Klaczko, 1999; Imasheva et al., 1999; Karan et al., 2000; Bubliy et al., 2001; Bubliy and Loeschcke, 2002; Debat et al., 2003, 2009; Shingleton et al., 2009; Peck and Maddrell, 2005). Previous work has demonstrated that some of the plastic response is aligned with clinal variation for a number of traits, including wing size and shape (Gilchrist et al., 2000; Pitchers et al., 2013; Klepsatel et al., 2014; Fabian et al., 2015). In addition to its role in flight performance (Ray et al., 2016), various aspects of wing morphology including size (Ewing, 1964; Abbott et al., 2010), shape (Menezes et al., 2013; Abbott et al., 2010), wing interference pattern (Katayama et al., 2014) and even wing musculature (Tracy et al., 2020) appear to be direct or indirect targets of sexual selection.

Evidence suggests that *Drosophila melanogaster* likely originates from the Miombo and Mopane forests in Zambia and Zimbabwe (Pool et al., 2012; Sprengelmeyer et al., 2020; Mansourian et al., 2018) and expanded its ancestral range in the past 13000 years, colonizing highland environments in Ethiopia between 2340 - 3060 years ago (Sprengelmeyer et al., 2020). In this study, we examine two populations from these altitudinally varying ranges. The low altitude (LA) population comes from Zambia and is assumed to be representative of ancestral *Drosophila* variation. The high altitude (HA) population comes from the highlands of Ethiopia. As is common with small insects adapting to life at high altitudes (reviewed in Hodkinson, 2005), a number of morphological, physiological and life history traits have evolved in the HA population (Pitchers et al., 2013; Pool et al., 2012; Lack et al., 2016a,b; Klepsatel et al., 2014; Fabian et al., 2015; Klepsatel et al., 2013). In particular, body size and wing size (and wing loading) have increased *∼*20% relative to the LA counterparts (Pitchers et al., 2013; Pool et al., 2012; Lack et al., 2016a,b; Klepsatel et al., 2014; Fabian et al., 2015; Klepsatel et al., 2013; Pesevski and Dworkin, 2020). Interestingly, there is evidence of concordance between the direction of temperature-induced plasticity and wing shape between lowland and highland populations (Pitchers et al., 2013). In this study, our goals were to (1) investigate how selection on wing form during adaptation to life at high altitude impacted plasticity (and more specifically, condition dependence) for sexual size and shape dimorphism, and (2) determine the relative contribution of shape-size allometries on the shape changes that were observed. We discuss our results within the context of the evolution of sexual shape dimorphism and the influence of allometry.

## Materials and Methods

### Drosophila husbandry

*Drosophila melanogaster* strains used in these experiments are a subset previously used in Lack et al. (2016a,b) and Pesevski and Dworkin (2020). Strains from the high altitude (HA) population were collected in Fiche, Ethiopia (3070m asl) in December 2011, while strains from the low altitude (LA) population were collected in Siavonga, Zambia (530m asl) (Supplementary Tables S1 and S2). There is no evidence that these two populations have had recent gene flow (Lack et al., 2016a,b). Prior to experimental treatments described in the following section, flies were raised on high protein food (HP; 1.5:1 protein to carbohydrate ratio) for two generations at room temperature to minimize maternal effects. Given that the strains derived from each population were maintained in the lab, we confirmed that our results were consistent with previous results and across experiments performed previously, as detailed in Pesevski and Dworkin.

### Experimental Design

#### Egg collections and experimental treatments

Adults were placed in egg laying chambers with apple juice agar plates with killed yeast patches (to avoid yeast growth and control for amount of food). After 12h egg laying windows, eggs were collected and placed in vials with high protein (HP) food, 50 eggs per vial (food is 2:1 carbohydrate to protein ratio). We performed two different treatment manipulation (1) varying food quality and (2) varying rearing temperature. Based on previous unpublished experiments in the lab, food treatments were done by raising flies on high quality (100% HP food), low quality (dilution to 15% of HP food) and very low quality food (dilution to 5% of HP food). While the viability for the 15% food was *∼*15%, for the 5% treatment it was very low (*∼*1%). As such, we decided to compare the 100% and 15% food treatments only. Flies under varying food treatments were reared at 24°. Flies for temperature treatments were raised flies only on high quality food (100%), in incubators at 18°C, 24°C and 28°C with a 12:12 light/dark cycles. Some results of the temperature manipulation were including in a previous study (Pesevski and Dworkin, 2020), but were done concurrently with the food manipulation study. For all temperature and 100% food treatments, 3 replicate vials of 50 eggs each were collected for each strain. For the 15% food treatment, 6 replicate vials were collected for each strain. Adults were collected after cuticle sceleratization and stored in 70% ethanol. This experiment was repeated twice (in January 2017 and June 2017), once with all of the lines and a second time with a subset of the lines to increase sample sizes for strains with low viability, and additional lines to assess block effects. The two replicate experiments are considered as two blocks in the analysis. Sample sizes for each strain and population are outlined in Supplementary Tables S1 and S2. We measured larval mass at the 3rd instar wandering stage to confirm that the nutritional manipulation reduced size and condition. To do this we repeated the egg laying experiment with a subset of strains. We added 25 eggs onto small petri dishes (30mm x 15mm) filled with either 100% or 15% food. We let the flies develop to the 3rd instar larval stage, determined their sex, washed, dried andd weighed them by pooling 5 individuals of the same sex and nutritional manipulation together for each independent biological sample.

#### Dissections and Imaging

A maximum of 20 male and 20 female adult flies were collected from each treatment/strain/replicate S1 and S2. The right wing of each fly was mounted on glass slides in 70% glycerol in PBS solution. Each wing was imaged using an Olympus DP30B camera mounted on an Olympus BX51 microscope (Olympus software version 3,1,1208) using a 4X objective (total of 40X magnification) and images were taken using cellSens Standard (version 1.14) software at 4060 x 3072 resolution.

#### Phenotyping

Wing size and shape were phenotyped using a modified version of the *“WINGMACHINE”* pipeline (Houle et al., 2003; Pitchers et al., 2019). Initial landmarks were placed at the humeral break on the leading edge of the wing and the alula notch on the trailing edge of the wing *tpsDig2* software (version 2.16). *“Wings”* software (version 3.7) was used to fit B splines to wing veins and wing margins and the splines were manually adjusted when necessary. The *x* and *y* coordinates of 12 landmarks and 36 semi-landmarks were extracted after Procrustes superimposition (removing position, orientation and scale) as well as the centroid size of each wing using *CPReader* software (version 1.12r). Approximate positions of landmarks and semi-landmarks are represented in Figure 2A.

### Analysis

Analyses were performed in *R* (v4.0.3) (R Core Team, 2018) using *R Studio* (v1.4.1103) on a *MacBook Pro*, running *macOS Big Sur* (v11.2.1). Linear mixed models were run using *lmer* from the *lme4* package (v1.1.26)(Bates et al., 2015), *glmmTMB* from the *glmmTMB* package (v1.0.2.1) (Magnusson et al., 2017) and *procD.lm* from the *geomorph* package (v3.3.2) (Adams et al., 2018). Contrasts and effects were compared using *emmeans* from the *emmeans* package (1.5.4) (Russell et al., 2018). We used *pairwise* from the *RRPP* package (v0.6.2) to perform pairwise comparisons of distance and directions of mean shapes between different groups (Collyer and Adams, 2020)

#### Modeling wing size and shape

Linear mixed models were fit with population, sex, environmental manipulation (either nutrition or temperature), and their interactions as fixed effects (up to 3rd order). We allowed intercept, sex effects and environmental treatment to vary as random effects of strain (line). We fit these models with both *lmer* and *glmmTMB* and confirmed estimates for parameters of interest were similar with both unconstrained and diagonal covariance structures for random effects.

Given the high-dimensional nature of our wing shape data, we used an alternative strategy for model fitting. Using *procD.lm* we fit models with population, sex and environmental manipulation and their interactions as fixed effects. To account for random effects of strain, we nested strain within population, and updated calculations of relevant F-ratios accordingly for terms in the model. ANOVA tables for both wing size and wing shape analyses are included in the supplementary data (Supplementary Tables S3, S4, S5 and S6)

#### Calculating and comparing SSD and SShD

We used a standard size dimorphism index (SDI) to calculate sexual size dimorphism (SSD), 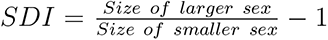. This SDI is generally used in species with female-biased SSD (Lovich and Gibbons, 1992; Fairbairn, 2007)). Mean female and mean male size were estimated for each strain and used to compare changes in SSD within and between populations and environmental manipulations (nutrition and temperature).

We assessed both the magnitude and direction of sexual shape dimorphism vectors. We calculated Procrustes distances between mean female and mean male shape vectors, and examined the direction of sexual shape change by calculating vector correlations (*r*) between female-male difference vectors among populations and environmental treatments (food and temperature). We generated confidence intervals for these statistics using non-parametric bootstraps (percentile intervals), sampling within groups.

#### Comparing allometric vectors

In our study design there are multiple sources contributing to size variation, which in turn influence shape-size allometric relationships. There is variation in size due to sex, genetic effects due to both population of origin, and strain (within population), plastic effects due to the experimental manipulations of temperature and food during development, and finally, any residual size variation among individuals. Given that the focus of this study is on the evolutionary changes in sexual shape dimorphism, we explored only a subset of these contributions in this study. We estimated shape-size allometric relationships per treatment group (i.e. partitioned by population, sex and either nutrition or temperature manipulation). Using *pairwise()* in *RRPP* we calculated vector correlations between allometry vectors, and the magnitudes of allometry vectors (i.e. shape change per unit change in size). To assess differences among groups for allometry, we fit reduced models assuming common allometry, and used *RRPP* to generate permutations of estimates under the full (group specific allometries and reduced (common allometry) models. An important limitation of the approach we took is that we did not account for strain specific allometries (strains within each population) in these models as we observed computational difficulties in model fitting when trying to account for these effects in *geomorph/RRPP*. As such the population effect represents a complete pooling (with respect to strain) within the focal groups under consideration. This is relevant and an important caveat as the work presented here, and in previous studies (Pitchers et al., 2013) have demonstrated strain-level variation in SShD.

#### Partitioning allometric, non-allometric and total SShD

To assess the relative contribution of changes in total SShD (*SShD_T_*) due to allometric (*SShD_A_*) and non-allometric (*SShD_NA_*) effects we used an approach related (but not identical) to Gidaszewski et al. (2009), assuming common allometry for males and females, and partitioned the relevant components. We estimated non-allometric SShD by computing Procrustes distance between the sex-specific “intercepts” in common allometry models. For allometric component of SShD we estimated the Procrustes distance between predicted vectors of shape variables at mean sizes of males and females respectively (but using the same “intercept” for both sexes) from this model. There are important caveats to this approach as it assumes a common allometric relationship between males and females. While these vectors are often similar in direction and magnitude (Tables 1 and 2), they are not identical, which will result in the partitioned values being difficult to interpret when these differ (see Gidaszewski et al. (2009) for further examples and discussion below). However, as an approximation and with appropriate caution, these still may provide some context to the relative contribution of allometric and non-allometric components to SShD. Importantly, the approach we took to estimate *SShD_NA_* and *SShD_A_* differ from some published studies (Sztepanacz and Houle, 2021; Gidaszewski et al., 2009) which computes one of the two components, and then uses the difference between these estimates and *SShD_T_* to compute the other value. As we show in the results and elaborate on in the discussion, when the assumption of common allometry is violated, this may produce estimates that are difficult to reconcile.

**Table 1:**
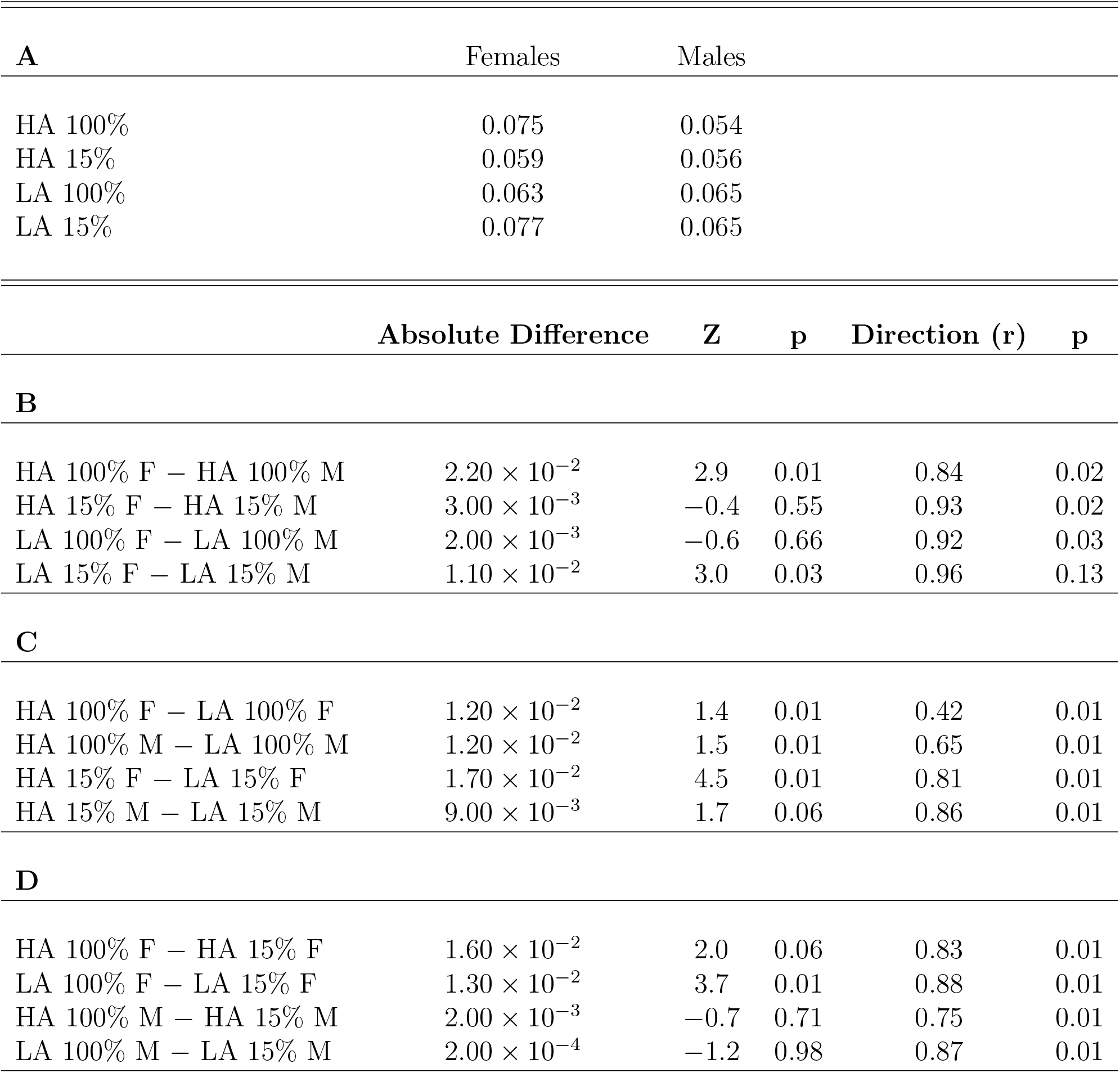
Pairwise comparisons of allometric vectors between populations and sexes for the food quality treatments. (A) Magnitudes (L2 norm) of allometry vectors for each group, and comparisons of allomeric vectors between (B) sexes, (C) populations and (D) food treatments. Comparisons for distance are calculated as absolute difference between the magnitudes of allometric vectors and comparisons for direction are calculated as vector correlation (*r*) <u>between the allometric vectors for each group.</u>

| **A** | Females | Males |
| --- | --- | --- |
| HA 100% | 0.075 | 0.054 |
| HA 15% | 0.059 | 0.056 |
| LA 100% | 0.063 | 0.065 |
| LA 15% | 0.077 | 0.065 |

|  | **Absolute Difference** | **Z** | **p** | **Direction (r)** | **p** |
| --- | --- | --- | --- | --- | --- |
| **B** |  |  |  |  |  |
| HA 100% F – HA 100% M | $2.20 \times 10^{-2}$ | 2.9 | 0.01 | 0.84 | 0.02 |
| HA 15% F – HA 15% M | $3.00 \times 10^{-3}$ | -0.4 | 0.55 | 0.93 | 0.02 |
| LA 100% F – LA 100% M | $2.00 \times 10^{-3}$ | -0.6 | 0.66 | 0.92 | 0.03 |
| LA 15% F – LA 15% M | $1.10 \times 10^{-2}$ | 3.0 | 0.03 | 0.96 | 0.13 |
| **C** |  |  |  |  |  |
| HA 100% F – LA 100% F | $1.20 \times 10^{-2}$ | 1.4 | 0.01 | 0.42 | 0.01 |
| HA 100% M – LA 100% M | $1.20 \times 10^{-2}$ | 1.5 | 0.01 | 0.65 | 0.01 |
| HA 15% F – LA 15% F | $1.70 \times 10^{-2}$ | 4.5 | 0.01 | 0.81 | 0.01 |
| HA 15% M – LA 15% M | $9.00 \times 10^{-3}$ | 1.7 | 0.06 | 0.86 | 0.01 |
| **D** |  |  |  |  |  |
| HA 100% F – HA 15% F | $1.60 \times 10^{-2}$ | 2.0 | 0.06 | 0.83 | 0.01 |
| LA 100% F – LA 15% F | $1.30 \times 10^{-2}$ | 3.7 | 0.01 | 0.88 | 0.01 |
| HA 100% M – HA 15% M | $2.00 \times 10^{-3}$ | -0.7 | 0.71 | 0.75 | 0.01 |
| LA 100% M – LA 15% M | $2.00 \times 10^{-4}$ | -1.2 | 0.98 | 0.87 | 0.01 |

**Table 2:** Pairwise comparisons of allometric vectors between populations and sexes for the temperature treatments. (A) Magnitudes (L2 norm) of allometric vectors for each group and comparisons of allomeric vectors between (B) sexes, (C) populations and (D) temperature treatments. Comparisons for distance are calculated as absolute difference between the magnitudes of allometric vectors and comparisons for direction are calculated as correlations (*r*) between allometric vectors for each group. (Values for 24°C are equivalent to 100% treatments in Table 1 and are therefore not represented here to avoid redundancy)

| **A** | Females | Males |
| --- | --- | --- |
| HA 18°C | 0.055 | 0.044 |
| HA 28°C | 0.056 | 0.057 |
| LA 18°C | 0.075 | 0.063 |
| LA 28°C | 0.070 | 0.072 |

|  | **Absolute Difference** | **Z** | **p** | **Direction (r)** | **p** |
| --- | --- | --- | --- | --- | --- |
| **B** |  |  |  |  |  |
| HA 18°C F – HA 18°C M | $1.11 \times 10^{-2}$ | 2.05 | 0.05 | 0.90 | 0.22 |
| HA 28°C F – HA 28°C M | $1.76 \times 10^{-3}$ | −0.96 | 0.80 | 0.51 | 0.01 |
| LA 18°C F – LA 18°C M | $1.15 \times 10^{-2}$ | 2.09 | 0.04 | 0.93 | 0.60 |
| LA 28°C F – LA 28°C M | $2.20 \times 10^{-3}$ | −0.49 | 0.57 | 0.96 | 0.92 |
| **C** |  |  |  |  |  |
| HA 18°C F – LA 18°C F | $1.93 \times 10^{-2}$ | 3.80 | 0.01 | −0.18 | 0.01 |
| HA 18°C M – LA 18°C M | $1.89 \times 10^{-2}$ | 3.90 | 0.01 | 0.17 | 0.01 |
| HA 28°C F – LA 28°C F | $1.46 \times 10^{-2}$ | 2.94 | 0.02 | 0.46 | 0.01 |
| HA 28°C M – LA 28°C M | $1.51 \times 10^{-2}$ | 2.21 | 0.05 | −0.04 | 0.01 |
| **D** |  |  |  |  |  |
| HA 18°C F – HA 24°C F | $2.00 \times 10^{-2}$ | 2.97 | 0.01 | 0.52 | 0.01 |
| HA 18°C M – HA 24°C M | $9.22 \times 10^{-3}$ | 0.78 | 0.17 | 0.50 | 0.01 |
| HA 18°C F – HA 28°C F | $1.62 \times 10^{-4}$ | −1.43 | 1.00 | 0.43 | 0.01 |
| HA 18°C M – LA 28°C M | $1.29 \times 10^{-2}$ | 1.39 | 0.13 | 0.85 | 0.17 |
| HA 24°C F – HA 28°C F | $1.95 \times 10^{-2}$ | 2.44 | 0.04 | 0.71 | 0.04 |
| HA 24°C M – HA 28°C M | $7.98 \times 10^{-3}$ | 0.46 | 0.22 | 0.51 | 0.02 |
| LA 18°C F – LA 24°C F | $1.15 \times 10^{-2}$ | 2.09 | 0.03 | 0.76 | 0.01 |
| LA 18°C M – LA 24°C M | $2.07 \times 10^{-3}$ | −0.8 | 0.75 | 0.76 | 0.01 |
| LA 18°C F – LA 28°C F | $4.55 \times 10^{-3}$ | 0.03 | 0.43 | 0.89 | 0.01 |
| LA 18°C M – LA 28°C M | $9.20 \times 10^{-3}$ | 1.02 | 0.15 | 0.90 | 0.33 |
| LA 24°C F – LA 28°C F | $7.00 \times 10^{-3}$ | 0.85 | 0.20 | 0.86 | 0.03 |
| LA 24°C M – LA 28°C M | $7.12 \times 10^{-3}$ | 0.61 | 0.23 | 0.85 | 0.14 |

## Results

### Wing size varies due to adaptive divergence and environmental plasticity, but SSD remains similar between populations

Consistent with previous observations (Pitchers et al., 2013; Klepsatel et al., 2014; Lack et al., 2016a,b; Pesevski and Dworkin, 2020), wing size of individuals from the high-altitude (HA) population are larger than those of the low-altitude (LA) population (Figure 1C). As expected, female-biased SSD was observed, with female wings being *∼*18% larger than males in both populations. Food and temperature treatments also affected wing size. Wing size in HA females raised on 100% food is *∼*17% larger than wing size in HA females raised at 15% food, while in HA males raised at 100% food it is 12% larger than in HA males raised at 15% food. Wing size in LA females raised at 100% food is 13% larger than wing size in LA females raised at 15% food, and in LA males raised at 100% food it is 9% larger than in LA males raised at 15% food (Figure 1A, Supplementary Table S3). Despite these substantial differences in wing size, changes in SSD between populations is proportional and not significantly different (Supplementary Table S3). Consistent with expectations of condition dependence, SSD is reduced due to the reduction in food quality (significant sex-by-diet term, Supplementary Table S3), but the reduction remains unchanged due to adaptive divergence between the HA and LA populations (sex-by-diet-by-population term is not significant, Supplementary Table S3, Figure 3C). In the HA population SSD is 0.136 in flies raised at 100% food, and it is reduced to 0.098 in flies raised at 15% food. Similarly, in the low altitude population, SSD is 0.134 in flies raised at 100% food and it is reduced to 0.106 in flies raised at 15% food (Supplementary Table S3).

**Figure 1:**
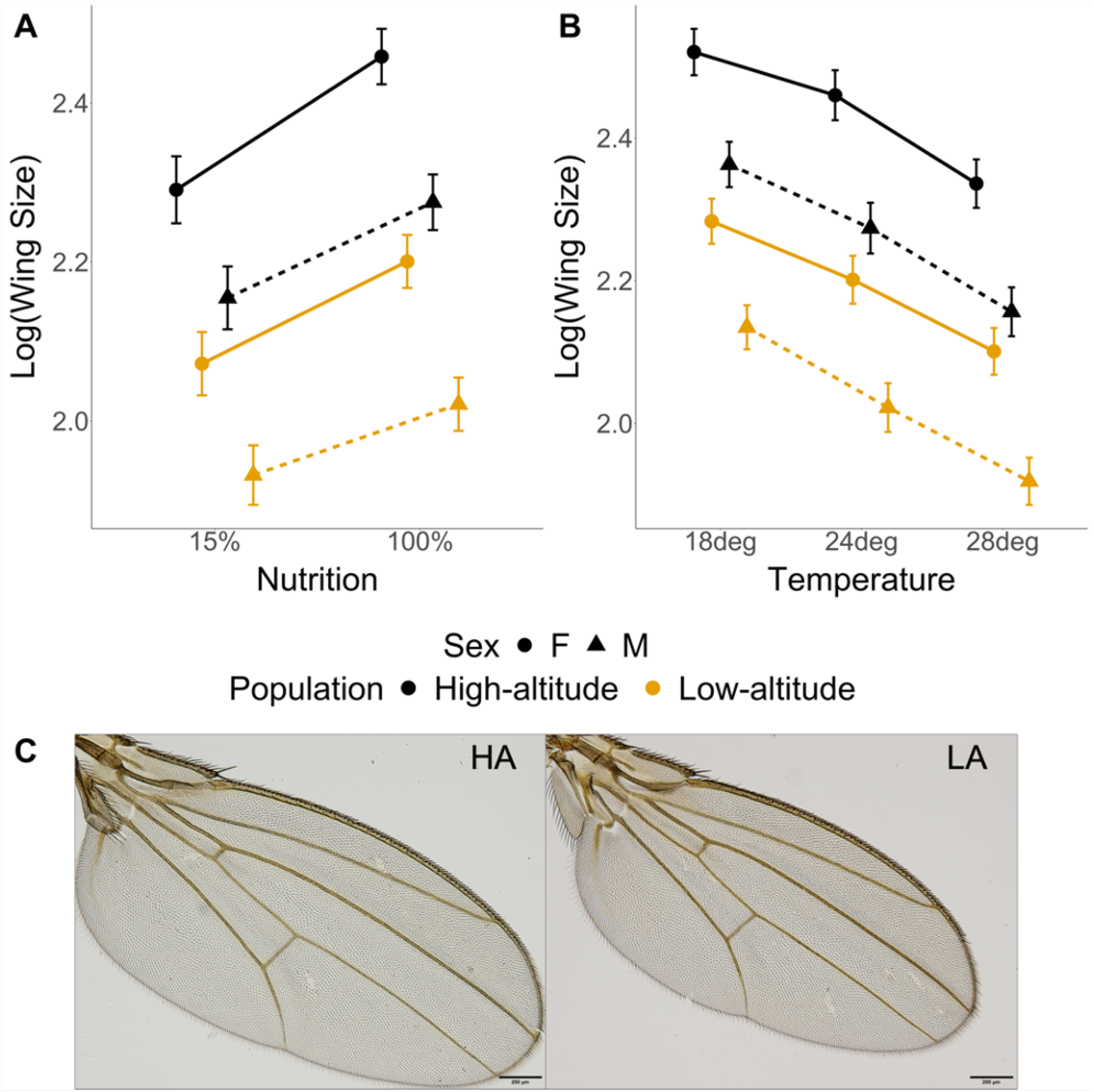
Wing size plasticity in high-altitude (HA) and low-altitude (LA) populations. A) Wing size plasticity in response to food quality is similar the HA and LA populations, despite substantial differences in overall sizes across both sex and population of origin. B) Wing size plasticity in response to rearing temperature in the HA and LA populations is similar, with an overall decrease in wing size with increase in temperature. C) Representative wings from the HA and LA populations. Estimates are averaged across strains within each population. Error bars represent 95% CIs

We measured larval mass at the wandering 3rd instar larval stage to determine if condition is being affected by food quality (Figure 2). We observed similar patterns for larval mass compared to patterns for wing size due to population, sex and nutrition (Figure 2). Importantly, we observed population and sex specific reduction in larva mass due to nutrition (Figure 2). The HA female larva were most sensitive to size reductions when reared on 15% food compared to other groups (Figure 2). This hints at strong population specific condition dependence of SSD for larval weight and potentially in overall body size.

**Figure 2:**
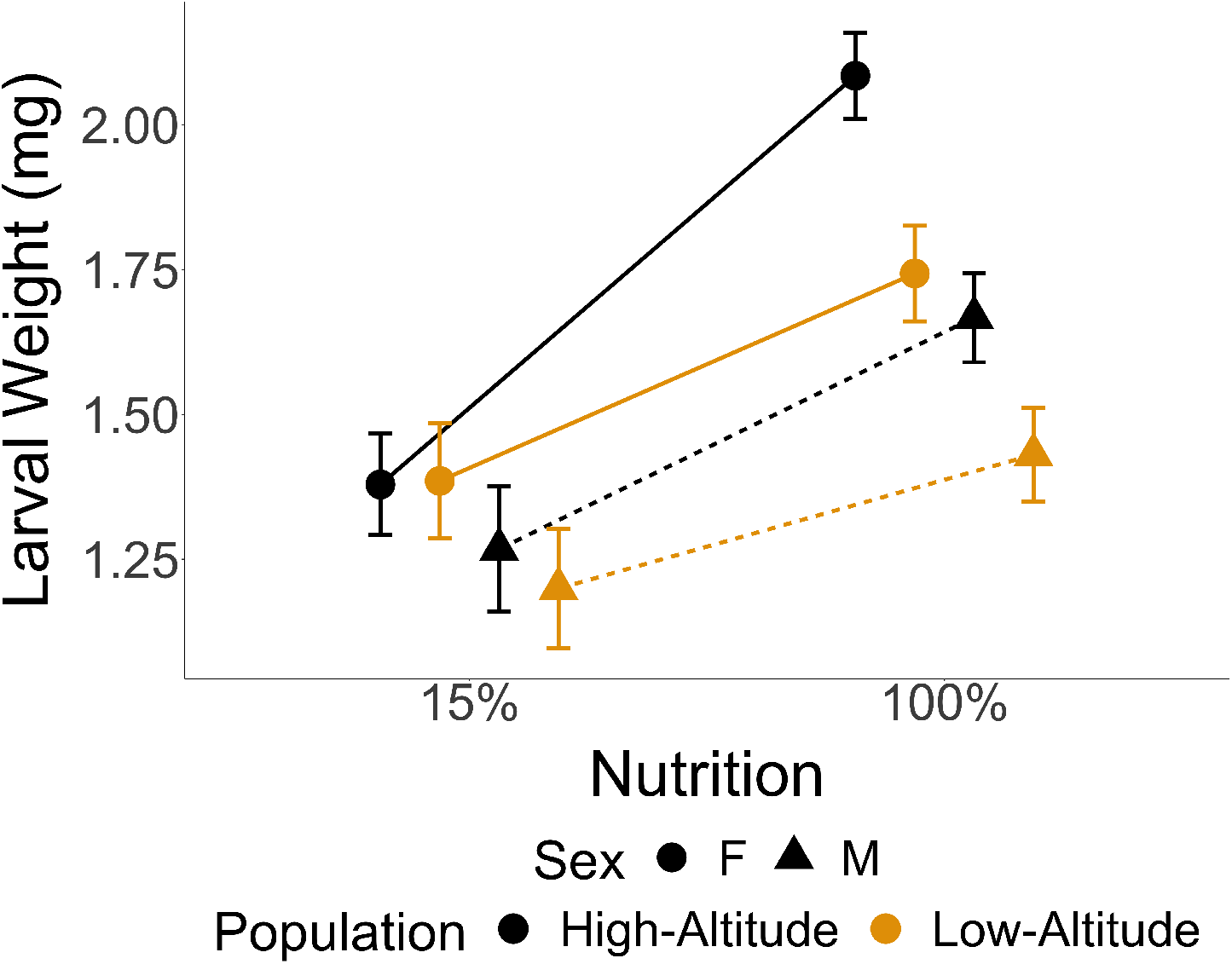
Larval weight plasticity in response to food quality in the high-altitude (HA) and low altitude (LA) populations. Larval weight in the HA females is the most sensitive to food quality reduction compared to the other groups.

Consistent with previous observations (Bitner–Mathe and Klaczko, 1999; David et al., 2009; Debat et al., 2003; Pesevski and Dworkin, 2020; Pitchers et al., 2013), an inverse relationship is seen between wing size and temperature (Figure 1B), and an overall increase in SSD with increasing temperature (Figure 3D). We observed evidence of sex-specific temperature plasticity as well as evidence for population-specific plasticity (significant sex-by-temperature and population-by-temperature interaction effects, Supplementary table S4, Figure 1B, Figure 3D). Modest population-specific changes in SSD in response to temperature (significant sex-by-population-by-temperature interaction effect, Supplementary Table S4, Figure 3D) was observed. In the HA population SSD is 0.117 at 18°C, 0.136 at 24°C and 0.138 at 28°C, while in the LA population SSD is 0.118 at 18°C, 0.134 at 24°C and 0.138 at 28°C (Figure 3D).

**Figure 3:**
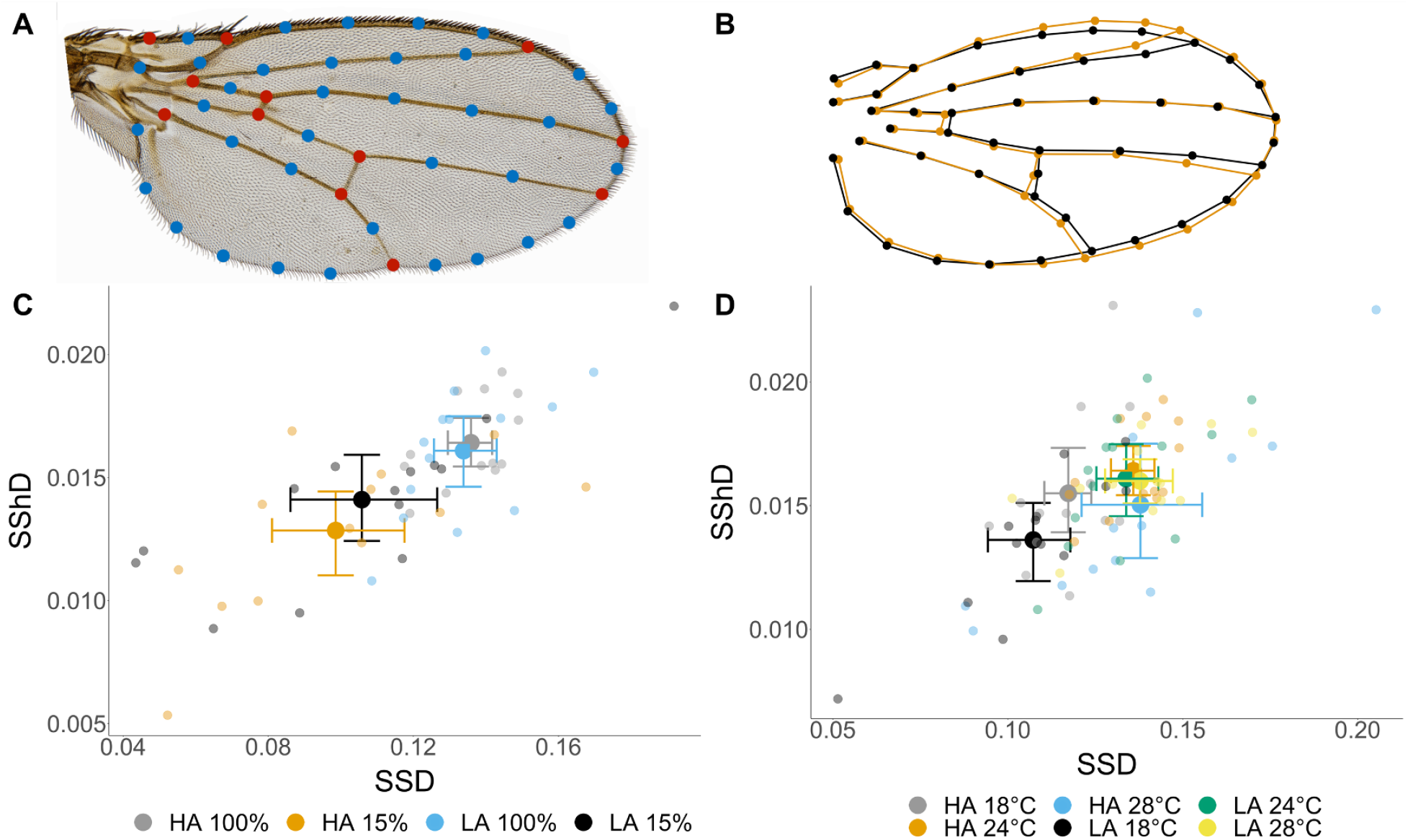
SShD variation in high-altitude (HA) and low-altitude (LA) populations. A) Landmarks (red) and semi-landmarks (blue) used in geometric morphometrics analysis to evaluate wing shape. B) Overlaid mean wing shapes of males (black) and females (orange) wings in the HA population at 24°C raised on 100% food quality as an example of SShD. C) Variation in SSD and SShD due to population and food quality. There is a greater reduction for SShD, but not for SSD, in the HA population compared to the LA population D) Variation in SSD and SShD due to population and rearing temperature. SShD is greatest at 24°in both HA and LA populations, but there is a greater reduction for SShD at 18°in the HA population compared to the LA population. For C and D, large opaque dots represent population means and small translucent dots represent means for each strain. Error bars represent 95% CIs

### Wing shape and total SShD vary due to environmental plasticity in a population-specific manner

Patterns of SShD between HA and LA populations were consistent with previous observations (Pitchers et al., 2013; Pesevski and Dworkin, 2020) (Figure 3, Supplementary Tables S5 and S6). We did not observe substantial population-specific difference in SShD, despite evidence for genetic variation in shape and sexual dimorphism among strains within populations (Supplementary Tables S5 and S6). Food quality had significant effects on wing shape, with some evidence of sex and population-specific differences (significant sex-by-diet and population-by-diet effects, Supplementary Table S5). We observed a reduction in SShD due to food quality, with a slightly greater effect in the HA population compared to the LA population (Figure 3C, Supplementary Table S5). In the HA population, SShD is 0.016 on 100% food which was reduced to *∼*0.013 on 15% food while in the LA population SShD is 0.016 on 100% food and it got reduced to *∼*0.014.

We observed consistent evidence of shape plasticity due to rearing temperature, including significant population-by-temperature and sex-by-temperature interactions (Supplementary Table S6). We observe slightly different patterns of SShD in response to temperature in HA and LA populations, where the greatest SShD is at 24°C in both populations, but the two populations differ greatly in the amount of SShD at 18°C (Figure 3D). For the HA population SShD is 0.015 at 18°C, 0.016 at 24°C and 0.015 at 28°C, and in the LA population SShD is 0.013 at 18°C, 0.016 at 24°C and 0.016 at 28°C.

Despite observed differences in magnitude of SShD due to food quality and rearing temperature in HA and LA populations, we see relatively similar changes in direction of SShD across populations and environmental treatments. Correlations among SShD vectors (across populations and environmental treatments) are all greater than 0.8, with many above 0.9 (Figure 4A,B). In particular, correlations of SShD vectors across populations are quite high, with somewhat lower values due to the food quality and temperature treatments (Figure 4A,B). This is consistent with much of the change in SShD being due to changes in magnitude, not direction for SShD.

**Figure 4:**
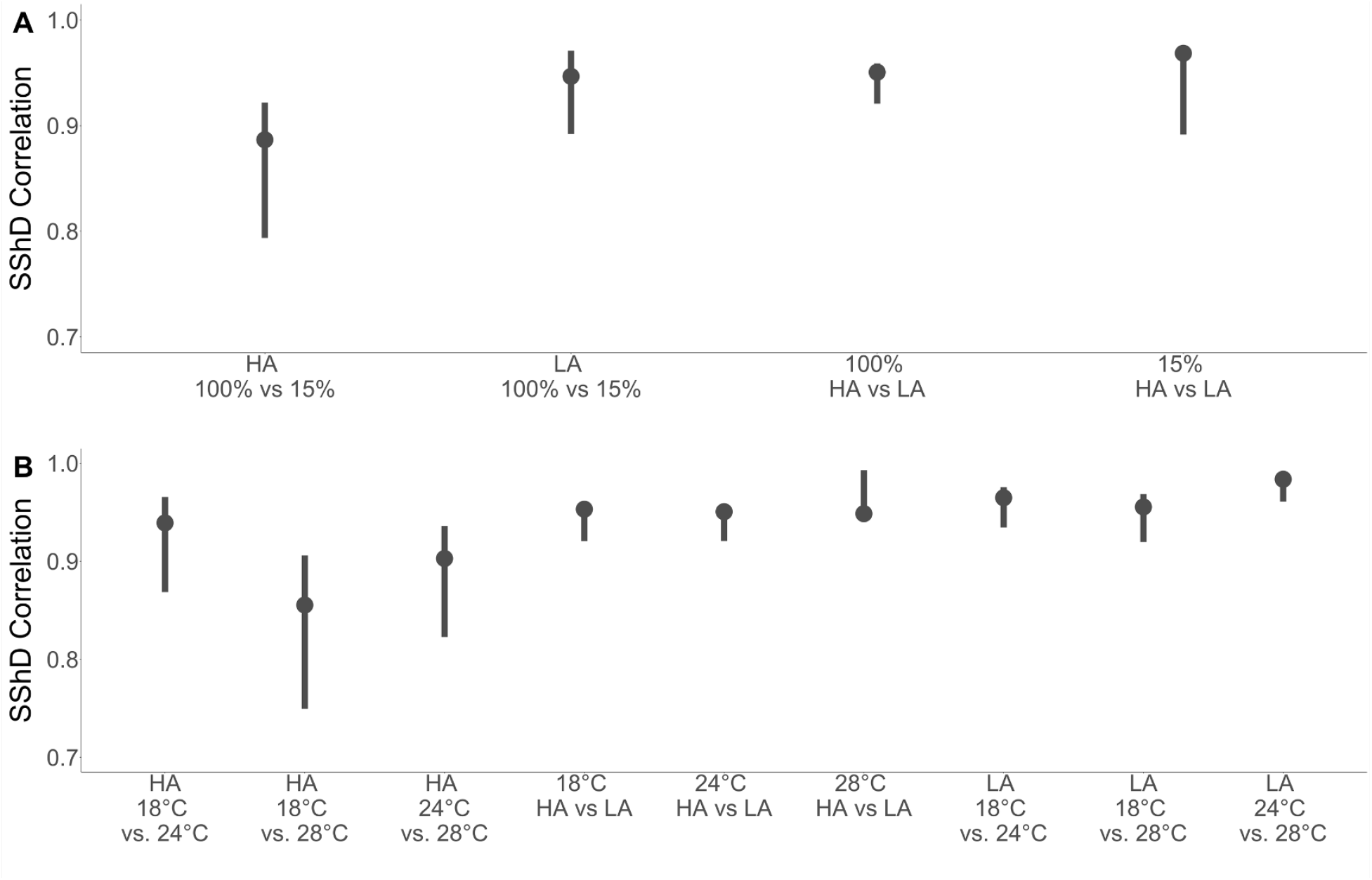
Correlations between SShD vectors in the high-altitude (HA) and low-altitude (LA) populations due to (A) food quality and (B) temperature plasticity. SShD vectors correlations are very high, demonstrating that variation in SShD is primarily in magnitude rather than direction. Error bars are bootstrap 95% CIs

### Differences in shape-size allometry are modest for SShD and environmental plasticity, but greater among diverged populations

We compared allometric vectors between populations, sex and environmental treatments, examining differences in magnitude (absolute difference) and direction (*r*) for allometric vectors between all groups (Tables 1 and 2). Overall, there are both sex-by-size and sex-by-population-by-size interactions (Supplementary tables S5 and S6) resulting from changes in magnitude, direction or both. Magnitude of shape change (per unit change in size) is modest, but consistently greater in females in both populations and across food quality treatments.

However, in direct paired comparisons across the sexes, differences are rarely significantly different from one another (Table 1A, Figure 5). The impact of food quality on magnitude of shape change is idiosyncratic, with modest evidence of changes only in LA females (Table 1A). This can be contrasted to the allometric effects across populations, where differences in magnitudes occur, but with the sign of differences reversing across sexes (Table 1A, Figure 5).

**Figure 5:**
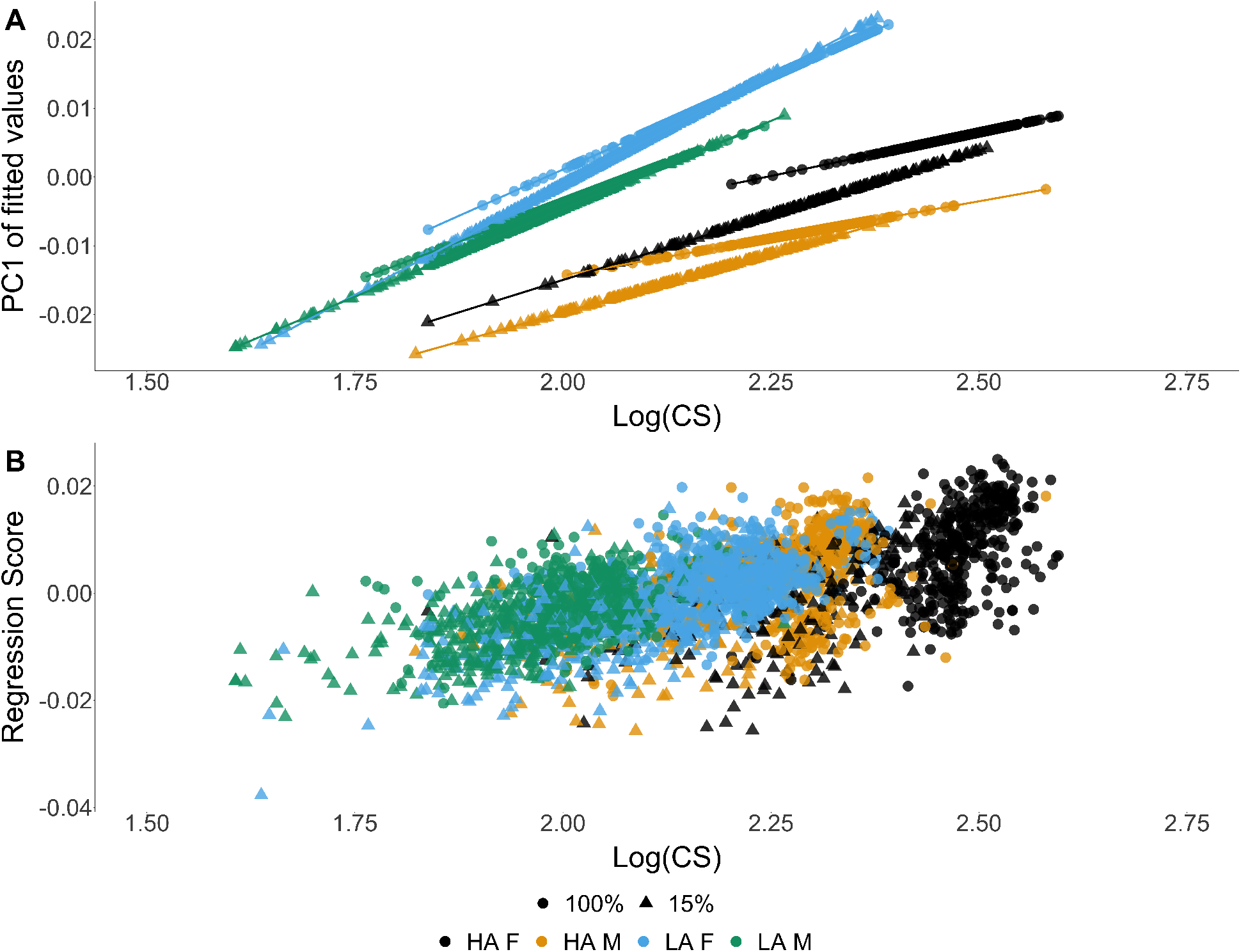
Size - shape allometry of wings in the high-altitude (HA) and low-altitude (LA) populations under different food treatments. (A) PC1 of fitted values plotted against wing size to visually represent the allometric vectors. (B) Regression shape scores plotted against wing size to visually represent the within groups for allometry

There is evidence of significant difference in the direction of allometric effects across sexes in both HA and LA populations, with the exception of the 15% food treatment in the LA population (Table 1B). In all of these cases, correlations among allometric vectors is high (Table 1B). While somewhat lower in the 15% food treatments, the direction of allometric effects remain similar (albeit significantly different), all with correlations of 0.84 and greater (Table 1B). This can be contrasted with greater differences between the allometric vectors in the two populations, with correlations closer to 0.5 in the 100% food treatments, but higher correlations (near 0.8) in the 15% food treatment (Table 1C). When comparing allometry vectors between food treatments (100% vs. 15%), we observe modest differences in both absolute distance and correlations, these differences being intermediate between the effects for sex and population (Table 1D).

The shape-size allometric patterns with respect to temperature treatments were quite distinct and more variable than for the food quality treatments (Table 2, Figure 6). Significant interactions between size and other factors in the model are observed (Supplemental table S6). Sex differences in magnitudes of allometry vectors due to rearing temperature remain modest, with little change to direction of allometric effects (Table 2A). There is a notable exception at 28°C (Table 2A). When comparing temperature effects within population and sex, we see a much more variable distribution of allometric effects, but consistent effects among populations (as observed for the population effects with varying food quality) in direction and magnitude (Table 2B-D, Figure 5). Allometric relationships also vary more across temperature manipulations within sex and population (Figure 6), contrasting with the allometric relationships under varying food quality treatments (Figure 5, Table 1).

**Figure 6:**
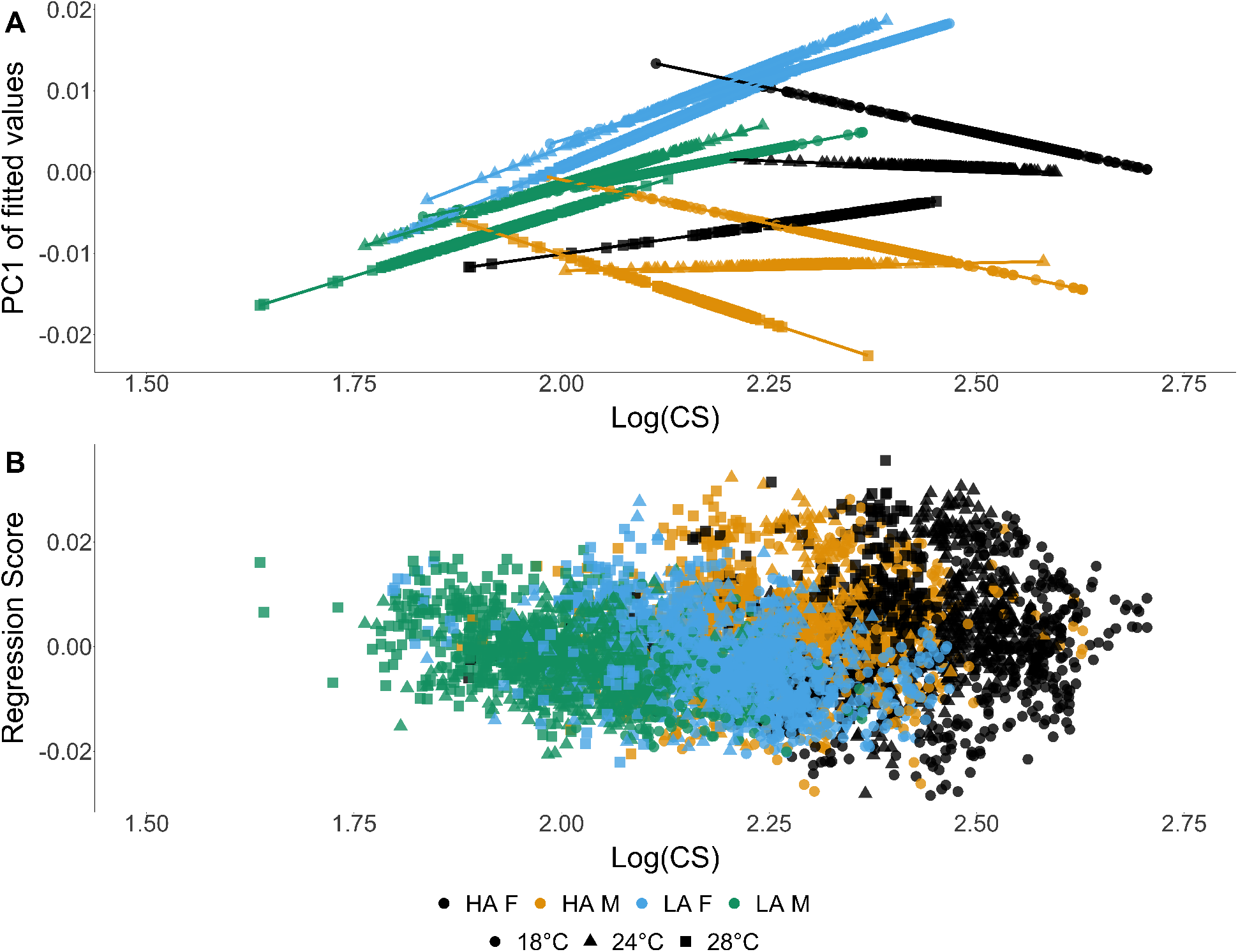
Size-shape allometry of wings in the high-altitude (HA) and low-altitude (LA) under different temperature treatments. (A) PC1 of fitted values plotted against wing size to visually represent the allometric vectors. (B) Regression shape scores plotted against wing size to visually represent the within groups for allometry

### Both allometric and non-allometric effects contribute to SShD

When considering the influence of size on shape, a common goal is to assess contributions of allometric and non-allometric effects. This is especially useful when studying SShD, because when there is SSD this may result in allometric SShD (*SShD_A_*) that can contribute a significant portion to total SShD (*SShD_T_*). The portion of *SShD_T_* that is not due to allometry is called non-allometric SShD (*SShD_NA_*) Most current approaches require the assumption of common allometry be at least approximately met (see (Klingenberg, 2016)). However, even when vector correlations are high, subtle violations of this assumption can result in difficult to interpret patterns (Gidaszewski et al., 2009). Despite these caveats, we assumed that since allometric vectors between the sexes had high correlations, we could assume common allometry and partitioned *SShD_T_* into *SShD_A_* and *SShD_NA_* components. Similar to previous results (Gidaszewski et al., 2009; Sztepanacz and Houle, 2021) we observed that *SShD_A_* and *SShD_NA_* contributed substantially to *SShD_T_* (Figure 7). However, we observe evidence that the assumption of common allometry is violated as values for *SShD_A_* and *SShD_NA_* do not sum to *SShD_T_*, and result in different estimates depending on how the components are calculated (Supplementary Table S10).

**Figure 7:**
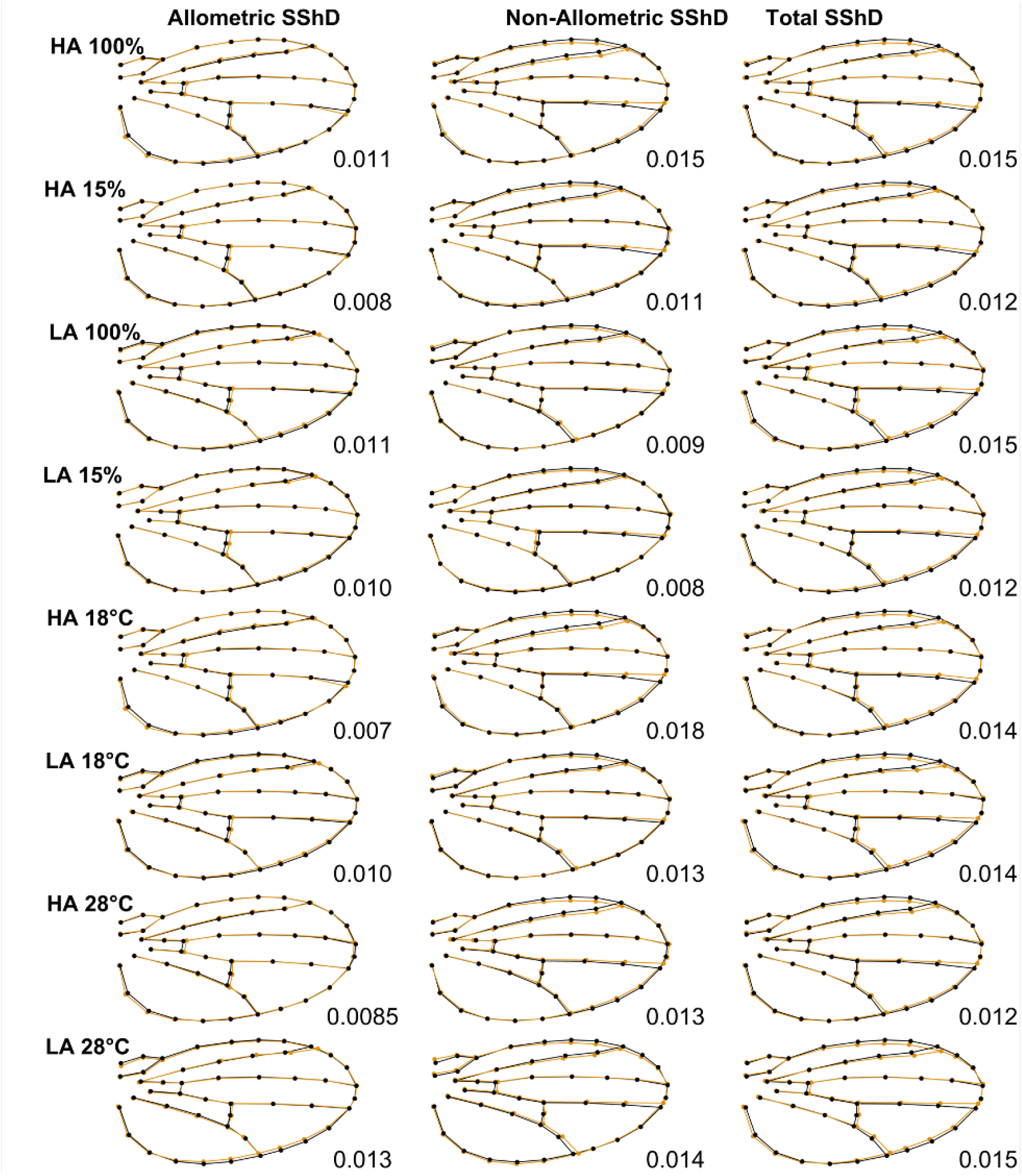
Allometric (*SShD_A_*), non-allometric (*SShD_NA_*) and total SShD (*SShD_T_*) within each population and environmental treatment. SShD values are calculated as the Procrustes distance (PD) between mean female and mean male shape. Both *SShD_A_* and *SShD_NA_* contribute towards *SShD_T_*, despite evidence that common allometry assumption between the sexes is being violated

## Discussion

### Influence of rapid adaptation on sex-specific plasticity

One of the main goals of this study was to examine how adaptive evolution influences sexspecific plasticity and condition dependence. As is typical for small insects (Hodkinson, 2005; Dillon et al., 2006), a number of traits contribute to the adaptive response to life at high altitudes, including body size, wing size and shape (Hodkinson, 2005; Pitchers et al., 2013; Pool et al., 2012; Lack et al., 2016a,b; Klepsatel et al., 2014, 2013; Fabian et al., 2015). The HA population used in this study has experienced strong directional selection for larger body size and disproportionately larger wing size (Pitchers et al., 2013; Pool et al., 2012; Lack et al., 2016a,b; Klepsatel et al., 2014; Fabian et al., 2015; Klepsatel et al., 2013). Wing shape has also evolved compared to the LA population, although the degree to which it is a direct target of selection is unclear (Pitchers et al., 2013). Aspects of wing morphology, including size and shape are targets of sexual selection in *D. melanogaster* (Ewing, 1964; Abbott et al., 2010; Menezes et al., 2013; Abbott et al., 2010), show sexual dimorphism within *D. melanogaster* and among other *Drosophila* species (Gidaszewski et al., 2009) and demonstrate sex-specific condition dependence. As such, we expected that the selection on size and shape contributing to the adaptive changes observed in the HA population may have altered genomic architecture such that it would influence sex-specific response to environmental variation (Connallon, 2015; Connallon et al., 2018). Contrary to these predictions, and despite substantial population differences in wing size and shape (Figures 1 and 3, Supplementary Tables S3,S4, S5, S6), changes in SSD and SShD are modest between populations (Figure 3C and D). As expected, we observed a reduction in SSD and SShD as a result of poor food quality (Figures 1A, 3C, Supplementary tables S3, S5). However, despite the evolved increase in overall size in the HA population, we did not observe a substantial increased sensitivity to food quality for SSD of wing size, and only a modest increase in sensitivity for SShD in the HA population (Figure 3C). Interestingly we observed a much stronger change in sex specific plasticity for larval mass, a proxy measure for body size. Larval mass showed a much greater reduction in SSD (sex specific condition dependence) in the high altitude population where females had a stronger response to food availability compared to the other groups (Figure 2). While we observed an increase of SSD and SShD as a result of increasing rearing temperature, the sensitivity to temperature of SSD and SShD in both populations was similar ((Figures 1B, 3D, Supplementary tables S4, S6)). This is similar to what has been observed for *r_fm_* number of traits for *Drosophila* populations along a latitudinal cline (Lasne et al., 2018). One possible explanation is that additional selective forces are maintaining the relative plastic response for wing morphology (Pesevski and Dworkin, 2020), but this may not be the case for overall body size. However, it is likely that a major target of selection in the HA population is wing loading (Klepsatel et al., 2014), and variation in wing shape can influence flight performance in lab environments for *Drosophila* species (Ray et al., 2016; Fraimout et al., 2018). Thus one hypothesis is that wing size and shape are under strong stabilizing selection in each environment resulting in greater environmental canalization.

### Evolution, plasticity and sexual dimorphism for shape—size allometry

The second goal of this study was to evaluate how much of the shape changes that were observed (across sex, population and rearing environments) was a consequence of allometry with size. This is a required step to ultimately identify whether wing shape was a direct target of selection resulting in the evolutionary wing shape changes observed in the HA population, or whether these changes are simply a correlated response due to allometry with size. Mean wing shape not only differs due to adaptive differences in the two populations, but there are significant mean shape differences due to sex, as well as due to nutritional and temperature plasticity (Supplementary Tables S5 and S6). However, as with most shape changes, shape — size allometry contributes a substantial amount to shape variation (Klingenberg, 2016), including in *Drosophila* (Gilchrist et al., 2000; Debat et al., 2003; Gidaszewski et al., 2009; Bolstad et al., 2015). Thus, we wanted to assess the relative contribution of allometric (and non-allometric) effects on wing shape. Given that we observed substantial differences in patterns in wing size variation due to population, sex and environmental plasticity (nutrition and rearing temperature) as well as the interaction among these, partitioning allometric effects can be challenging (Figure 1, Supplementary tables S3, S4, S5, S6). We utilized multiple approaches to asses the contribution of allometric and non-allometric effects on wing shape because of the complicated nature of this endeavour. In addition to discussing this in the context of our results, we also briefly discuss the effectiveness of each approach in the context of understanding allometric effects. As is often the case, partitioning the effects using a multivariate linear model (and allowing size effects to vary according to sex and other predictors) suggests that we observe “significant” variation in allometric relationships across groups (Supplementary Tables S5 and S6). Given the relatively large sample sizes used in this study, it is not surprising we detect statistically significant effects. More importantly, we examined the relative contribution of changes in direction and magnitude of shape change associated with changes in size, as summarized in Tables 1 and 2, and Figures 5 and 6 Despite significant interactions between sex and size, changes in magnitude of allometry (i.e magnitude of shape change per unit change in size) is generally modest as are changes in direction of allometric effects (Table 1, Figure 5). This pattern seems to largely hold with respect to temperature mediated effects on sexual dimorphism for allometry (Table 2, Figure 6). This can be contrasted with differences in allometric effects among populations which show evidence of divergence for direction and magnitude, often showing a surprising lack of concordance in direction. Whether this reflects that shape itself or shape-size allometry has been a direct target of selection is unclear. One clear finding from our study is how different environmental mediators of change in size (and shape) like nutrition and rearing temperature, elicit different patterns of allometry in terms of population of origin and sexual dimorphism (contrast Figure 5 and 6). In particular, while allometric effects in response to food quality elicit reasonably aligned changes in shape across sexes and populations, influence of rearing temperature differs substantially. An analogous pattern has been observed for *D. melanogaster* for multivariate allometry of size when both rearing temperature and food availability were varied (Shingleton et al., 2009). The degree to which these differences reflect potentially adaptive temperature plasticity is unclear, but previous work demonstrated alignment between shape changes between HA and LA populations and temperature plasticity (Pitchers et al., 2013).

In this study, we attempted to infer relative contributions of allometric and non-allometric effects on SShD by partitioning these components. Current approaches require an assumption of common allometry between the sexes within each population and rearing condition (Sztepanacz and Houle, 2021), and violations of that assumption can result in misleading inferences. While this assumption was clearly violated across populations, allometric patterns across the sexes were generally the most similar with respect to both direction and magnitude (Tables 1B,2B, Figures 5A, 6A), as has been observed previously (Gidaszewski et al., 2009; Testa and Dworkin, 2016). Given the relatively modest violation of the assumption of common allometry, we explored this approach to determine the degree to which it might be informative. Despite having vector correlations among allometry vectors between mostly greater than 0.9 between males and females, we still observe problematic effects where summed contribution of *SShD_A_* and *SShD_NA_* are in excess of *SShD_T_* (Figure 7). To the extent that any meaningful inference can be made, our evidence suggests that the *SShD_A_* and *SShD_NA_* components are relatively evenly contributing to *SShD_T_* (Figure 6). Very similar effects and likely similar problems have been observed when partitioning SShD among *Drosophila* species (Gidaszewski et al., 2009; Sztepanacz and Houle, 2021). Overall, this suggests that even when the direction of allometric effects appears very similar, partitioning allometric and non-allometric under an assumption of common allometry may be misleading. Thus the approach used for partitioning such effects, can result in substantially different answers (Supplementary Table S10). Care must still be taken when there is no significant interaction effect between sex and size, as this often is a function of studies with modest sample sizes (or where size variation is small) (Klingenberg, 2016). In such cases, allometric relationships can be estimated with high uncertainty, suggesting an apparent (but potentially incorrect) similarity in relationship. Developing approaches to partition allometric effects that are robust to such assumptions remain an important open problem in geometric morphometrics.

Despite being among the first studies examining the influence of rapid adaptation on condition dependent sexual dimorphism (and sex-specific plasticity in general) in a moderately female biased SSD system, our study is not without limitations. First, we are studying the interplay of adaptation, sexual dimorphism and plasticity within a single pair of adaptively diverged populations. Even though we did not observe evidence of changes in patterns of sex-specific plasticity as a consequence of rapid adaptive evolution, it does not rule out such changes in other populations and/or species, particularly in systems where adaptive and sexual selection forces may be misaligned. While sexual selection is known to generally operate on body size (Testa et al., 2013), wing size (Ewing, 1964; Abbott et al., 2010), shape (Menezes et al., 2013; Abbott et al., 2010) and other aspects of wing morphology (Katayama et al., 2014) of *D. melanogaster*, we do not know the strength or direction of sexual selection in the populations we examined. Natural selection for greater body and wing size in the HA population could have altered the strength of sexual selection in the wings of HA population. We observed that the degree of SSD for the wing is similar in the two populations hinting at potentially unchanged patterns of sexual selection in the two populations, but that remains to be confirmed experimentally.

As with any manipulative experiment, differential viability among treatments may result in biased sampling of individuals used for phenotyping, despite efforts for random sampling. In this study, we measured egg-to-adult viability (Figures S1 and S2) observing more than a *∼*65% reduction in viability for individuals raised on 15% food (survivorship at about 11– 16%) relative to flies raised on the 100% food (survivorship at about 45–46%). Viability in the low altitude population was significantly higher than for the HA population for the 15% food treatment (supported by a significant nutrition:population interaction, Supplementary Table S8). This suggests that the HA population fitness may be more strongly affected by poor food quality than the LA population. This ‘invisible fraction’ may represent a nonrandom subset of phenotypes as well. We did not observe any differences in viability between populations at different rearing temperatures (Supplementary Table S9).

This study focused on the evolutionary changes in wing morphology, and other than larval mass, we did not take measurements of other body parts. Previous work has demonstrated that the evolved increase in size was for a number of traits (Pitchers et al., 2013; Pool et al., 2012; Lack et al., 2016a,b; Klepsatel et al., 2014, 2013; Fabian et al., 2015) but that the increase in wing size was disproportionately large. However, it is known that different organs of *Drosophila* respond to the influence of nutritional and temperature manipulations to differing degrees (Shingleton et al., 2009). In retrospect, these measurements would have been very useful to explore the differences in plastic response for different traits and overall body size, and potentially scaling effects of overall body size and the wing. Despite this, the strength of our study is the exploration of wing shape and SShD, as well as, the allometric and non-allometric contributions of shape variation in these two populations. The results from the examination of larval mass suggest that we may be underestimating the condition dependence of SSD by focusing only on the wing because we observe a heightened condition dependence of SSD for larval weight in the HA population, particularly in the HA females (Supplementary Figure 1, supported by significant sex:population:nutrition effect, Supplementary table S7).

Finally, strains used in this study were established in 2011, while the experimental manipulations (performed simultaneously) were done in 2018. As such, potential issues due to inbreeding, genetic drift and to a lesser extent lab domestication can occur. As multiple strains were used from each population (Supplementary tables S1 and S2), the impact of drift on allelic frequencies across all strains within a population will be in generally modest. Lab domestication does not appear to have had a substantial impact on morphological changes. In a previous study we examined phenotypic variation of these same strains at two different time points (*∼*5 years apart) and observed they were highly correlated, and remained consistent with previous studies about adaptive divergence for these populations (Pesevski and Dworkin, 2020). This suggests minimal influence of lab domestication on traits under study. This is perhaps not surprising as when established as strains from single females *N_e_* is small for each strain, and thus selection is unlikely to be particularly efficient except for variants with large effects on fitness.

In summary, although we observed sex-specific plasticity for wing size, we did not observe major changes in these patterns as a result of adaptive divergence. Interestingly, we observed slight increase in sex-specific plasticity for wing shape. We observed that differences in wing shape due to sex, adaptive divergence and plasticity are a product of both allometric and non-allometric components. Despite the limitations of the method for partitioning allometric and non-allometric SShD, we observe that both allometric and non-allometric components of SShD contribute to overall SShD. These findings are the beginning of the exploration of the interplay between adaptive evolution, sexual dimorphism and sex-specific plasticity (Ceballos and Valenzuela, 2011). However, our understating of these patterns remains limited and further theoretical and empirical work is needed to explore these relationships further. Although recently there has been a greater interest in examining the coevolution of sexual dimorphism and condition dependence (Bonduriansky, 2007b; Rohner and Blanckenhorn, 2018; Cotton et al., 2004a; Stillwell and Fox, 2007), there is still a gap in understanding of the underlying mechanisms that govern the relationships between sexual dimorphism and condition dependence. Further examination of influence of local adaptation on the relationship between sexual dimorphism and condition dependence is also necessary because natural populations are often under multiple selective pressures at the same time. Although the HA population is an interesting natural example to examine the interplay between condition dependent sexual dimorphism and local adaptation, there is a substantial need to explorem this in different species, including a range of traits that vary in the magnitude and direction of SSD (and SShD), would me most welcome in order to fully understand these processes and how they interact.

## Acknowledgements

We wish to thank Dr. John Pool and Dr. Justin Lack for sending strains. We thank Sachin Davis, Melissa Rezik, Ravina Dhami, and Vikram Bhagavat for help with dissections and imaging. This work was funded by an NSERC (Canada) Discovery and Discovery accelerator grant to ID. We would like to thank NSERC and McMaster University for continued financial support during the covid-19 pandemic.

## Supplemental Figures

**Figure S1:**
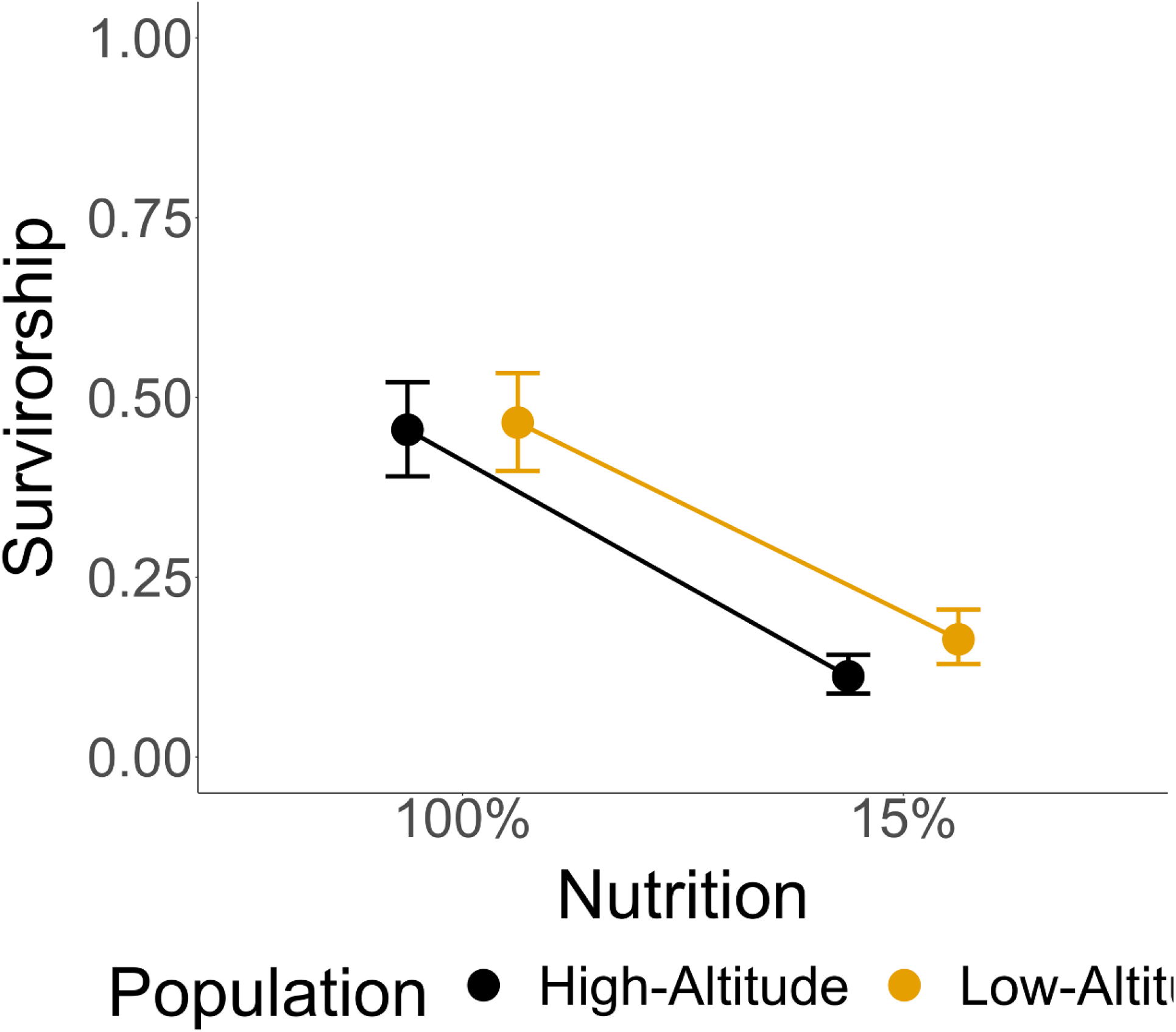
Survival Nutrition

**Figure S2:**
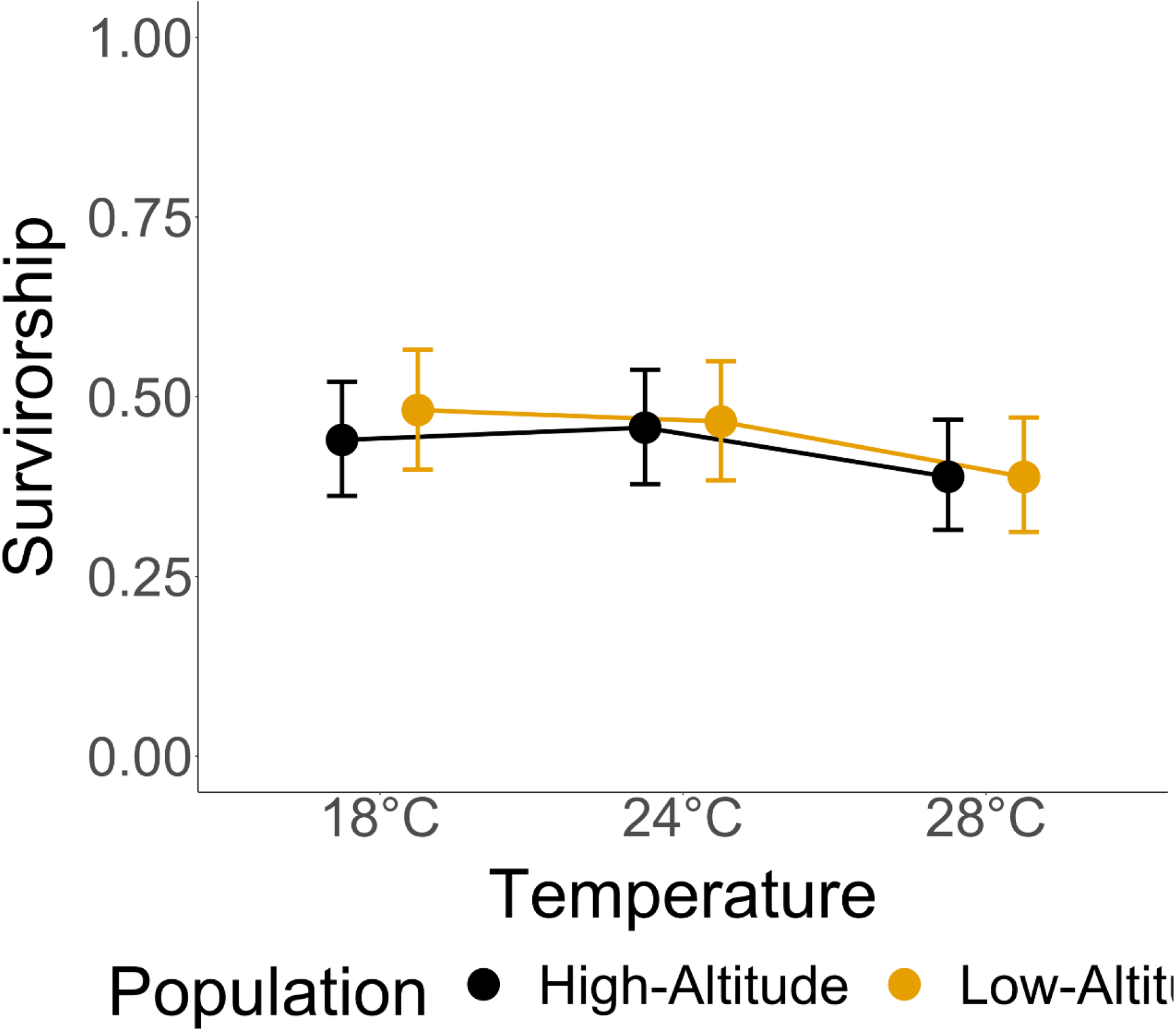
Survival Temperature

## Supplemental Tables

**Table S1:** Sample sizes of fly strains for Nutritional manipulation

| Line | Population | $N_{100\%F}$ | $N_{15\%F}$ | $N_{100\%F}$ | $N_{15\%M}$ |
| --- | --- | --- | --- | --- | --- |
| ef112 | High-Altitude | 10 | 13 | 23 | 9 |
| ef117 | High-Altitude | 37 | 5 | 27 | 5 |
| ef119 | High-Altitude | 24 | 13 | 9 | 19 |
| ef131 | High-Altitude | 27 | 22 | 39 | 19 |
| ef136 | High-Altitude | 42 | 10 | 40 | 2 |
| ef15 | High-Altitude | 41 | 2 | 47 | 6 |
| ef19 | High-Altitude | 12 | 6 | 13 | 6 |
| ef39 | High-Altitude | 48 | 59 | 58 | 56 |
| ef59 | High-Altitude | 7 | 6 | 8 | 12 |
| ef65 | High-Altitude | 28 | 5 | 25 | 4 |
| ef73 | High-Altitude | 43 | 18 | 63 | 23 |
| ef16 | High-Altitude | 27 | 25 | 26 | 24 |
| ef98 | High-Altitude | 16 | 18 | 20 | 12 |
| **Total** | **High-Altitude** | **362** | **202** | **398** | **197** |
| zi124 | Low-Altitude | 56 | 23 | 43 | 24 |
| zi159 | Low-Altitude | 72 | 30 | 60 | 24 |
| zi160 | Low-Altitude | 22 | 29 | 19 | 10 |
| zi186 | Low-Altitude | 8 | 37 | 8 | 28 |
| zi216 | Low-Altitude | 47 | 17 | 56 | 10 |
| zi251 | Low-Altitude | 44 | 29 | 71 | 34 |
| zi254 | Low-Altitude | 71 | 48 | 81 | 46 |
| zi311 | Low-Altitude | 45 | 55 | 43 | 41 |
| zi357 | Low-Altitude | 31 | 14 | 33 | 9 |
| zi360 | Low-Altitude | 50 | 34 | 39 | 24 |
| zi366 | Low-Altitude | 33 | 39 | 28 | 22 |
| zi367 | Low-Altitude | 35 | 19 | 23 | 17 |
| zi455 | Low-Altitude | 32 | 16 | 37 | 22 |
| zi455 | Low-Altitude | 55 | 31 | 38 | 24 |
| **Total** | **Low-Altitude** | **601** | **421** | **579** | **335** |

**Table S2:** Sample sizes of fly strains for Temperature manipulation

| Line | Population | $N_{18^{\circ}CF}$ | $N_{18^{\circ}CM}$ | $N_{28^{\circ}CF}$ | $N_{28^{\circ}CM}$ |
| --- | --- | --- | --- | --- | --- |
| ef112 | High-Altitude | 28 | 18 | 22 | 9 |
| ef117 | High-Altitude | 25 | 41 | 23 | 27 |
| ef119 | High-Altitude | 20 | 16 | 11 | 9 |
| ef131 | High-Altitude | 26 | 35 | 41 | 27 |
| ef136 | High-Altitude | 31 | 45 | 31 | 32 |
| ef15 | High-Altitude | 33 | 32 | 49 | 40 |
| ef19 | High-Altitude | 7 | 6 | 9 | 10 |
| ef39 | High-Altitude | 34 | 48 | 44 | 52 |
| ef59 | High-Altitude | 15 | 10 | 5 | 3 |
| ef65 | High-Altitude | 21 | 17 | 6 | 8 |
| ef73 | High-Altitude | 65 | 61 | 35 | 32 |
| ef16 | High-Altitude | 42 | 36 | 43 | 30 |
| ef98 | High-Altitude | 25 | 19 | 13 | 8 |
| **Total** | **High-Altitude** | **372** | **374** | **632** | **287** |
| zi124 | Low-Altitude | 73 | 70 | 65 | 60 |
| zi159 | Low-Altitude | 41 | 39 | 42 | 31 |
| zi160 | Low-Altitude | 31 | 21 | 0 | 0 |
| zi186 | Low-Altitude | 26 | 37 | 21 | 23 |
| zi216 | Low-Altitude | 26 | 29 | 10 | 4 |
| zi251 | Low-Altitude | 37 | 38 | 38 | 40 |
| zi254 | Low-Altitude | 90 | 71 | 67 | 70 |
| zi311 | Low-Altitude | 47 | 29 | 44 | 32 |
| zi357 | Low-Altitude | 25 | 18 | 17 | 19 |
| zi360 | Low-Altitude | 36 | 49 | 40 | 35 |
| zi366 | Low-Altitude | 23 | 33 | 16 | 15 |
| zi367 | Low-Altitude | 8 | 8 | 23 | 10 |
| zi455 | Low-Altitude | 32 | 35 | 15 | 16 |
| zi461 | Low-Altitude | 0 | 0 | 37 | 18 |
| **Total** | **Low-Altitude** | **495** | **477** | **435** | **373** |

**Table S3:** ANOVA for wing size fitting sex, population and nutrition as fixed effects and line as random effect

|  | Chisq | Df | Pr(>Chisq) |
| --- | --- | --- | --- |
| Sex | $2.1 \times 10^3$ | 1 | $< 2.0 \times 10^{-16}$ |
| Pop | $1.2 \times 10^2$ | 1 | $< 2.0 \times 10^{-16}$ |
| Nut | $1.0 \times 10^2$ | 1 | $< 2.0 \times 10^{-16}$ |
| Sex:Pop | $1.6 \times 10^{-2}$ | 1 | $9.0 \times 10^{-1}$ |
| Sex:Nut | 76 | 1 | $< 2.0 \times 10^{-16}$ |
| Pop:Nut | 1.9 | 1 | $1.6 \times 10^{-1}$ |
| Sex:Pop:Nut | $5.8 \times 10^{-1}$ | 1 | 0.44 |

**Table S4:** ANOVA for wing size fitting sex, population and temperature as fixed effects and line as random effect

|  | Chisq | Df | Pr(>Chisq) |
| --- | --- | --- | --- |
| Sex | $3.0 \times 10^3$ | 1 | $< 2.0 \times 10^{-16}$ |
| Pop | $1.1 \times 10^2$ | 1 | $< 2.0 \times 10^{-16}$ |
| Temp | $9.5 \times 10^2$ | 2 | $< 2.0 \times 10^{-16}$ |
| Sex:Pop | $7.0 \times 10^{-1}$ | 1 | $4.0 \times 10^{-1}$ |
| Sex:Temp | $8.5 \times 10^1$ | 2 | $< 2.0 \times 10^{-16}$ |
| Pop:Temp | 4.1 | 2 | $1.3 \times 10^{-1}$ |
| Sex:Pop:Temp | 2.5 | 2 | $2.9 \times 10^{-1}$ |

**Table S5:** MANOVA table for wing shape fitting size, sex, population and nutrition as fixed effects

| Effect | Df | SS | MS | Rsqr | F | Z | p (>F) |
| --- | --- | --- | --- | --- | --- | --- | --- |
| Size | 1 | $1.5 \times 10^{-1}$ | $1.5 \times 10^{-1}$ | $8.1 \times 10^{-2}$ | 4.9 | 2.9 | $3.0 \times 10^{-3}$ |
| Pop | 1 | $2.1 \times 10^{-1}$ | $2.1 \times 10^{-1}$ | $1.1 \times 10^{-1}$ | 6.7 | 3.5 | $< 1.0 \times 10^{-3}$ |
| Sex | 1 | $4.4 \times 10^{-2}$ | $4.4 \times 10^{-2}$ | $2.3 \times 10^{-2}$ | 39 | 6.3 | $< 1.0 \times 10^{-3}$ |
| Nut | 1 | $2.2 \times 10^{-2}$ | $2.2 \times 10^{-2}$ | $1.2 \times 10^{-2}$ | 33 | 6.3 | $< 1.0 \times 10^{-3}$ |
| Pop:Line | 25 | $7.8 \times 10^{-1}$ | $3.1 \times 10^{-2}$ | $4.1 \times 10^{-1}$ | $1.6 \times 10^2$ | 31 | $< 1.0 \times 10^{-3}$ |
| Size:Pop | 1 | $3.1 \times 10^{-3}$ | $3.1 \times 10^{-3}$ | $1.6 \times 10^{-3}$ | $1.0 \times 10^{-1}$ | -3.5 | 1.0 |
| Size:Sex | 1 | $1.8 \times 10^{-3}$ | $1.8 \times 10^{-3}$ | $9.5 \times 10^{-4}$ | 1.6 | 1.2 | $1.1 \times 10^{-1}$ |
| Pop:Sex | 1 | $8.3 \times 10^{-4}$ | $8.3 \times 10^{-4}$ | $4.4 \times 10^{-4}$ | $7.4 \times 10^{-1}$ | $-6.4 \times 10^{-1}$ | $7.2 \times 10^{-1}$ |
| Size:Nut | 1 | $7.0 \times 10^{-4}$ | $7.0 \times 10^{-4}$ | $3.7 \times 10^{-4}$ | 1.0 | $-6.7 \times 10^{-2}$ | $5.3 \times 10^{-1}$ |
| Pop:Nut | 1 | $2.1 \times 10^{-3}$ | $2.1 \times 10^{-3}$ | $1.1 \times 10^{-3}$ | 3.1 | 2.6 | $8.0 \times 10^{-3}$ |
| Sex:Nut | 1 | $2.7 \times 10^{-3}$ | $2.7 \times 10^{-3}$ | $1.4 \times 10^{-3}$ | 6.9 | 4.7 | $< 1.0 \times 10^{-3}$ |
| Size:Pop:Line | 25 | $2.6 \times 10^{-2}$ | $1.0 \times 10^{-3}$ | $1.4 \times 10^{-2}$ | 5.2 | $1.4 \times 10^1$ | $< 1.0 \times 10^{-3}$ |
| Pop:Sex:Line | 25 | $2.8 \times 10^{-2}$ | $1.1 \times 10^{-3}$ | $1.5 \times 10^{-2}$ | 5.7 | $1.6 \times 10^1$ | $< 1.0 \times 10^{-3}$ |
| Size:Pop:Sex | 1 | $1.2 \times 10^{-3}$ | $1.2 \times 10^{-3}$ | $6.6 \times 10^{-4}$ | 1.1 | $5.6 \times 10^{-1}$ | $2.9 \times 10^{-1}$ |
| Pop:Nut:Line | 25 | $1.7 \times 10^{-2}$ | $6.8 \times 10^{-4}$ | $8.9 \times 10^{-3}$ | 3.4 | $1.5 \times 10^1$ | $< 1.0 \times 10^{-3}$ |
| Size:Pop:Nut | 1 | $4.4 \times 10^{-4}$ | $4.4 \times 10^{-4}$ | $2.3 \times 10^{-4}$ | 1.1 | $3.7 \times 10^{-1}$ | $3.5 \times 10^{-1}$ |
| Pop:Sex:Nut | 1 | $7.2 \times 10^{-4}$ | $7.2 \times 10^{-4}$ | $3.8 \times 10^{-4}$ | 1.8 | 1.5 | $6.9 \times 10^{-2}$ |
| Size:Pop:Sex:Line | 25 | $7.3 \times 10^{-3}$ | $2.9 \times 10^{-4}$ | $3.8 \times 10^{-2}$ | 1.5 | 4.4 | $< 1.0 \times 10^{-3}$ |
| Size:Pop:Nut:Line | 25 | $7.8 \times 10^{-3}$ | $3.1 \times 10^{-4}$ | $4.1 \times 10^{-3}$ | 1.6 | 5.3 | $< 1.0 \times 10^{-3}$ |
| Pop:Sex:Nut:Line | 25 | $1.0 \times 10^{-3}$ | $4.0 \times 10^{-4}$ | $5.2 \times 10^{-3}$ | 2.0 | 7.7 | $< 1.0 \times 10^{-3}$ |
| Size:Pop:Sex:Nut | 1 | $5.0 \times 10^{-4}$ | $5.0 \times 10^{-4}$ | $2.6 \times 10^{-4}$ | 1.3 | $8.2 \times 10^{-1}$ | $2.0 \times 10^{-1}$ |
| Size:Pop:Sex:Nut:Line | 25 | $6.7 \times 10^{-3}$ | $2.7 \times 10^{-4}$ | $3.5 \times 10^{-3}$ | 1.3 | 3.2 | $2.0 \times 10^{-3}$ |
| Residuals | $2.9 \times 10^3$ | $5.7 \times 10^{-2}$ | $2.0 \times 10^{-4}$ | $3.0 \times 10^{-1}$ | | | |
| Total | $3.1 \times 10^3$ | 1.9 | | | | | |

**Table S6:** MANOVA table for wing shape fitting size, sex, population and temperature as fixed effects

| Effect | Df | SS | MS | Rsq | F | Z | p (>F) |
| --- | --- | --- | --- | --- | --- | --- | --- |
| Size | 1 | $1.2 \times 10^{-1}$ | $1.2 \times 10^{-1}$ | $3.8 \times 10^{-2}$ | 2.4 | 1.5 | $7.9 \times 10^{-2}$ |
| Pop | 1 | $3.0 \times 10^{-1}$ | $3.0 \times 10^{-1}$ | $9.6 \times 10^{-2}$ | 5.9 | 3.2 | $2.0 \times 10^{-3}$ |
| Sex | 1 | $1.2 \times 10^{-1}$ | $1.2 \times 10^{-1}$ | $3.8 \times 10^{-2}$ | 68 | 7.9 | $1.0 \times 10^{-3}$ |
| Temp | 2 | $1.2 \times 10^{-1}$ | $6.0 \times 10^{-2}$ | $3.8 \times 10^{-2}$ | 38 | 7.2 | $1.0 \times 10^{-3}$ |
| Pop:Line | 25 | 1.3 | $5.1 \times 10^{-2}$ | $4.0 \times 10^{-1}$ | $2.5 \times 10^2$ | 29 | $1.0 \times 10^{-3}$ |
| Size:Pop | 1 | $1.2 \times 10^{-2}$ | $1.2 \times 10^{-2}$ | $3.9 \times 10^{-3}$ | $2.4 \times 10^{-1}$ | -1.5 | $9.2 \times 10^{-1}$ |
| Size:Sex | 1 | $6.2 \times 10^{-3}$ | $6.2 \times 10^{-3}$ | $2.0 \times 10^{-3}$ | 3.5 | 2.9 | $2.0 \times 10^{-3}$ |
| Pop:Sex | 1 | $1.2 \times 10^{-3}$ | $1.2 \times 10^{-3}$ | $3.6 \times 10^{-4}$ | $6.4 \times 10^{-1}$ | $-9.3 \times 10^{-1}$ | $8.3 \times 10^{-1}$ |
| Size:Temp | 2 | $5.2 \times 10^{-3}$ | $2.6 \times 10^{-3}$ | $1.6 \times 10^{-3}$ | 1.7 | 1.5 | $6.7 \times 10^{-2}$ |
| Pop:Temp | 2 | $1.0 \times 10^{-2}$ | $5.0 \times 10^{-3}$ | $3.1 \times 10^{-3}$ | 3.2 | 3.7 | $1.0 \times 10^{-3}$ |
| Sex:Temp | 2 | $4.4 \times 10^{-3}$ | $2.2 \times 10^{-3}$ | $1.4 \times 10^{-3}$ | 4.6 | 5.0 | $1.0 \times 10^{-3}$ |
| Size:Pop:Line | 25 | $5.5 \times 10^{-2}$ | $2.2 \times 10^{-3}$ | $1.7 \times 10^{-2}$ | 11 | 28 | $1.0 \times 10^{-3}$ |
| Pop:Sex:Line | 25 | $4.5 \times 10^{-2}$ | $1.8 \times 10^{-3}$ | $1.4 \times 10^{-2}$ | 9.0 | 21 | $1.0 \times 10^{-3}$ |
| Size:Pop:Sex | 1 | $1.1 \times 10^{-3}$ | $1.1 \times 10^{-3}$ | $3.5 \times 10^{-4}$ | $6.2 \times 10^{-1}$ | $-8.2 \times 10^{-1}$ | $8.0 \times 10^{-1}$ |
| Pop:Temp:Line | 48 | $7.4 \times 10^{-2}$ | $1.5 \times 10^{-3}$ | $2.3 \times 10^{-2}$ | 7.8 | 21 | $1.0 \times 10^{-3}$ |
| Size:Pop:Temp | 2 | $1.7 \times 10^{-3}$ | $8.5 \times 10^{-4}$ | $5.4 \times 10^{-4}$ | $5.5 \times 10^{-1}$ | -1.6 | $9.5 \times 10^{-1}$ |
| Pop:Sex:Temp | 2 | $1.4 \times 10^{-3}$ | $6.9 \times 10^{-4}$ | $4.3 \times 10^{-4}$ | 1.4 | 3.3 | $2.0 \times 10^{-3}$ |
| Size:Pop:Sex:Line | 25 | $1.1 \times 10^{-2}$ | $4.5 \times 10^{-4}$ | $3.6 \times 10^{-3}$ | 2.3 | 8.5 | $1.0 \times 10^{-3}$ |
| Size:Pop:Temp:Line | 48 | $2.0 \times 10^{-2}$ | $4.3 \times 10^{-4}$ | $6.5 \times 10^{-3}$ | 2.1 | 14 | $1.0 \times 10^{-3}$ |
| Pop:Sex:Temp:Line | 48 | $2.3 \times 10^{-3}$ | $4.8 \times 10^{-4}$ | $7.3 \times 10^{-3}$ | 2.4 | 15 | $1.0 \times 10^{-3}$ |
| Size:Pop:Sex:Temp | 2 | $1.1 \times 10^{-3}$ | $5.3 \times 10^{-4}$ | $3.3 \times 10^{-4}$ | 1.1 | 1.1 | $3.3 \times 10^{-1}$ |
| Size:Pop:Sex:Temp:Line | 48 | $1.1 \times 10^{-2}$ | $2.4 \times 10^{-4}$ | $3.6 \times 10^{-3}$ | 1.2 | 3.1 | $1.0 \times 10^{-3}$ |
| Residuals | $4.8 \times 10^3$ | $9.4 \times 10^{-1}$ | $2.0 \times 10^{-4}$ | $3.0 \times 10^{-1}$ | | | |
| Total | $5.1 \times 10^3$ | 3.2 | | | | | |

**Table S7:** ANOVA for larval weight fitting block, sex, population and nutrition as fixed effects and line as random effect

|  | Chisq | Df | Pr(>Chisq) |
| --- | --- | --- | --- |
| Block | 4.8 | 1 | $2.8 \times 10^{-2}$ |
| Sex | $1.5 \times 10^2$ | 1 | $< 2.2 \times 10^{-16}$ |
| Pop | 24 | 1 | $1.2 \times 10^{-6}$ |
| Nut | $2.9 \times 10^2$ | 1 | $< 2.2 \times 10^{-16}$ |
| Sex:Pop | $8.9 \times 10^{-1}$ | 1 | $3.5 \times 10^{-1}$ |
| Sex:Nut | 17 | 1 | $3.7 \times 10^{-5}$ |
| Pop:Nut | 25 | 1 | $5.0 \times 10^{-7}$ |
| Sex:Pop:Nut | 2.8 | 1 | $9.3 \times 10^{-2}$ |

**Table S8:** Generalized linear mixed model for survivorship fitting population and nutrition as fixed effects and line as a random effect

|  | Chisq | Df | Pr(>Chisq) |
| --- | --- | --- | --- |
| Pop | 1.2 | 1 | $2.6 \times 10^{-1}$ |
| Nut | $2.2 \times 10^3$ | 1 | $< 2.2 \times 10^{-16}$ |
| Pop:Nut | 31 | 1 | $3.015 \times 10^{-8}$ |

**Table S9:** Generalized linear mixed model for survivorship fitting population and temperature as fixed effects and line as a random effect

|  | Chisq | Df | Pr(>Chisq) |
| --- | --- | --- | --- |
| Pop | $7.7 \times 10^{-2}$ | 1 | $7.8 \times 10^{-1}$ |
| Temp | 85 | 2 | $< 2 \times 10^{-16}$ |
| Pop:Temp | 5.7 | 2 | $5.7 \times 10^{-2}$ |

**Table S10:**
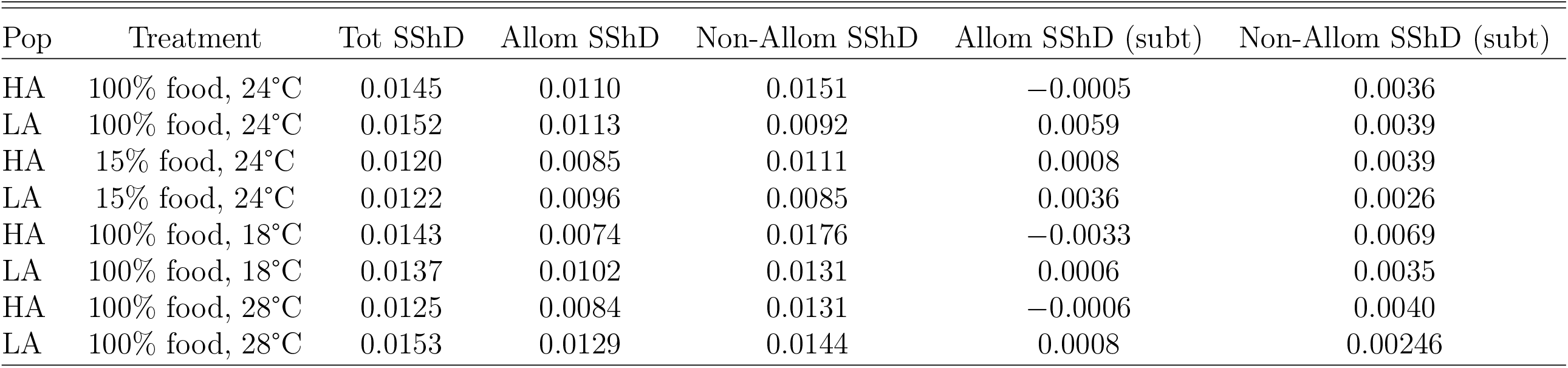
Calculating Partitioned SShD in two ways

| Pop | Treatment | Tot SShD | Allom SShD | Non-Allom SShD | Allom SShD (subt) | Non-Allom SShD (subt) |
| --- | --- | --- | --- | --- | --- | --- |
| HA | 100% food, 24°C | 0.0145 | 0.0110 | 0.0151 | −0.0005 | 0.0036 |
| LA | 100% food, 24°C | 0.0152 | 0.0113 | 0.0092 | 0.0059 | 0.0039 |
| HA | 15% food, 24°C | 0.0120 | 0.0085 | 0.0111 | 0.0008 | 0.0039 |
| LA | 15% food, 24°C | 0.0122 | 0.0096 | 0.0085 | 0.0036 | 0.0026 |
| HA | 100% food, 18°C | 0.0143 | 0.0074 | 0.0176 | −0.0033 | 0.0069 |
| LA | 100% food, 18°C | 0.0137 | 0.0102 | 0.0131 | 0.0006 | 0.0035 |
| HA | 100% food, 28°C | 0.0125 | 0.0084 | 0.0131 | −0.0006 | 0.0040 |
| LA | 100% food, 28°C | 0.0153 | 0.0129 | 0.0144 | 0.0008 | 0.00246 |

